# Nrf2 regulates the activation-driven expansion of CD4^+^ T-cells by differentially modulating glucose and glutamine metabolism

**DOI:** 10.1101/2024.04.18.590146

**Authors:** Aprajita Tripathi, Debolina Dasgupta, Anil Pant, Ashlyn Bugbee, Nanda Kumar Yellapu, Ben H. Y. Choi, Shailendra Giri, Kalyani Pyaram

**Author notes:** Correspondence, X: @ImmunolKalyani.

## Abstract

**SUMMARY:** Upon antigenic stimulation, CD4^+^T-cells undergo clonal expansion, elevating their bioenergetic demands and utilization of nutrients like glucose and glutamine. The nuclear factor erythroid 2-related factor 2 (Nrf2) is a well-known regulator of oxidative stress, but its involvement in modulating the metabolism of CD4^+^T-cells remains unexplored. Here, we elucidate the role of Nrf2 beyond the traditional antioxidation, in modulating activation-driven expansion of CD4^+^T-cells by influencing their nutrient metabolism. T-cell-specific activation of Nrf2 enhances early activation and IL-2 secretion, upregulates TCR-signaling, and increases activation-driven proliferation of CD4^+^T-cells. Mechanistically, high Nrf2 inhibits glucose metabolism through glycolysis but promotes glutamine metabolism via glutaminolysis to support increased T-cell proliferation. Further, Nrf2 expression is temporally regulated in activated CD4^+^T-cells with elevated expression during the early activation, but decreased expression thereafter. Overall, our findings uncover a novel role of Nrf2 as a metabolic modulator of CD4^+^T-cells, thus providing a framework for improving Nrf2-targeting therapies and T-cell immunotherapies.

## INTRODUCTION

CD4^+^T-cells or T-helper (Th) cells are key players of the adaptive immune system as they orchestrate and coordinate the immune responses against antigens^1–3^. The antigen presentation by major histocompatibility complex (MHC) class II molecules to the T-cell receptor (TCR) aided by the co-stimulatory signal such as CD28, initiates activation of naïve T-cells^3^. Once stimulated, T-cells undergo clonal expansion by several rounds of proliferation to produce effector T-cells for specific immune responses. It is coupled with metabolic programming to transition from a quiescent state reliant on oxidative phosphorylation (OXPHOS) to a highly glycolytic and anabolic state to produce ATP to meet the energy demands of rapid clonal expansion and cytokine functions^4–7^. They rely on glucose-fueled aerobic glycolysis and glycolysis-driven metabolic intermediates for growth and expansion^8^. Additionally, the amino acid Glutamine (Gln) is critical for the activation-driven proliferation of T-cells^9^ and for Th cell fate decision^10,11^. Although the role played by glucose and Gln metabolism in T-cell activation and effector functions is becoming clear, much remains to be understood about the upstream factors and signaling molecules governing them.

Activation triggers a rapid increase in the production and levels of reactive oxygen species (ROS) within T-cells by NADPH oxidases and mitochondrial respiratory complexes. Maintaining ROS homeostasis is vital for ensuring optimal T-cell responses^12–15^, as ROS-driven oxidative stress leads to defective T-cell viability and functions^16,17^. As such, redox balance in mammalian cells is tightly regulated by the trimeric antioxidation complex, comprising of Kelch-like ECH-associated protein 1 (Keap1), nuclear factor erythroid 2–related factor 2 (Nrf2) and cullin 3 (Cul3)^18,19^. Keap1 binds and represses transcription factor Nrf2 under homeostatic conditions via ubiquitin-proteasome pathway^18,20,21^. During oxidative stress, conformational changes in Keap1 dislodge Nrf2, which translocates into the nucleus and binds to the antioxidant response element (ARE) along with small musculoaponeurotic fibrosarcoma (Maf) proteins to regulate the transcription of antioxidant-responsive genes like NAD(P)H quinone oxidoreductase 1 (Nqo1) and heme oxygenase-1 (HO-1)^19,20^. Nrf2 has emerged as a pleiotropic transcription factor regulating metabolism, inflammation and autophagy^22^.

Despite its established role as a metabolic regulator in cancer cells^23,24^, whether Nrf2 influences T-cell metabolism remains largely unknown. In a first study, we reported that Nrf2 controls the development and effector functions of innate Natural Killer T (NKT) cells by regulating glucose uptake and mitochondrial biogenesis^25^. In patients with childhood rheumatism high Nrf2 was reported to correlate with increased metabolic state of T-cells^26^. A recent study in CD8^+^T-cells reported that Asparagine restriction activates Nrf2 and metabolically reprograms their anti-tumor responses^27^. It is still unclear if and how Nrf2 affects the activation and cellular metabolism of CD4^+^T-cells. Studies using Nrf2 activators like tert-butylhydroquinone, dimethyl fumarate, and CDDO-Im indicate an impact of Nrf2 on CD4^+^T-cell activation and proliferation^28,29,30^. However, there are several discrepancies and contradictory results from these inhibitor studies, likely due to their off-target effects and toxicity.

In this study, using T-cell-specific activation or deletion of Nrf2, we demonstrate that it differentially regulates nutrient dependency of CD4^+^T-cells to impact activation-driven expansion. High Nrf2 expression was found to enhance TCR signaling, increase proliferative capacity, and upregulate glutamine metabolism in activated CD4^+^T-cells whilst suppressing glycolysis.

## RESULTS

### High Nrf2 promotes activation-driven proliferation of CD4^+^ T-cells

The expression levels of Nrf2 were reported to increase post-activation in human T-cells ^31,32^ but its significance is unknown. We probed if Nrf2 modulates T-cell activation and promotes the expansion of CD4^+^T-cells. As in humans, we found that the protein expression of Nrf2 increases at days 1 and 2 of activation in mouse CD4^+^T-cells but, decreases by day 3 (Figure S1A). The subsequent decrease in Nrf2 expression correlated with increased Keap1 expression indicating that Nrf2 is regulated in a Keap1-dependent manner. Activated T-cells undergo clonal expansion as the first step of their effector functions^33^. Therefore, we investigated the role of Nrf2 in the expansion of activated CD4^+^T-cells. We used mice with T-cell specific deletion of *Keap1* gene (Figure S1B) (Keap1^fl/flCD4Cre^, referred to as Keap1-KO; K-KO) leading to constitutively high Nrf2 activity^32^ and expression in activated CD4^+^T-cells (Figure S1C). Naïve CD4 T-cells from WT and K-KO mice were isolated and activated using anti-CD3 and anti-CD28 antibodies *in vitro* (Figure 1A) to mimic T-cell stimulation. Interestingly, K-KO CD4^+^T-cells with high Nrf2 demonstrated a significant increase in proliferation compared to the WT CD4^+^T-cells upon activation, as measured by the dilution of cell trace dye (Figure 1B). Accordingly, K-KO CD4^+^T-cells also depicted an increase in cell size (Figures 1C and S1D). To concurrently study and compare the effects of low Nrf2, we included mice with T-cell-specific deletion of Nrf2 (Nrf2^fl/flCD4Cre^, called Nrf2-KO or N-KO) in our study. Compared to the WT CD4^+^T-cells, we noted a modest decrease in the proliferation of Nrf2-KO CD4^+^T-cells upon activation (Figure S1E). We then sought to validate these findings *in vivo* using BrdU (5-bromo-2’-deoxyuridine) after injecting the mice with anti-CD3 antibodies to activate T-cells (Figures 1D and S1F). K-KO mice had higher frequencies of BrdU^+^CD4^+^T-cells while Nrf2-KO mice had lower frequencies compared to the WT mice (Figure 1E). To establish that Nrf2 modulates antigen-specific expansion of CD4^+^T-cells, we generated Ova-specific TCR-transgenic (OTII)^34^ Keap1-KO and Nrf2-KO mice and studied *in vivo* BrdU incorporation following antigen challenge with ovalbumin (Figure 1F). Higher percentages of BrdU^+^CD4^+^T-cells were observed in OTII-Keap1-KO mice than in OTII-WT or OTII-Nrf2-KO mice (Figure 1G) indicating enhanced antigen-driven expansion of Nrf2-high CD4^+^T-cells. Similar results were observed with *in vitro* activation using Ova peptide (OVA_323-339_) (Figure 1H). Notably, despite their higher proliferative capacity, lower Keap1-KO CD4^+^T-cell counts were observed in the *in vitro* cultures compared to WT and Nrf2-KO cells (Fig. 1I) and thus we tested if the lower cell counts were due to increased cell death. Indeed, more apoptotic cells were detected among the activated Keap1-KO CD4^+^ T-cells (Figure 1J). Furthermore, Gene Ontology (GO) enrichment analysis of RNA sequencing data (Figure S1G) demonstrated an upregulation of genes involved in lymphocyte proliferation and homeostasis in Keap1-KO over WT CD4^+^T-cells (Figure S1H). We also noted upregulation of genes associated with programmed cell death and apoptosis in activated Keap1-KO CD4^+^T-cells (Fig. S1I). Taken together, these findings indicate that temporally regulated Nrf2 drives the proliferation of activated CD4^+^ T-cells and modulates their homeostasis.

**Figure 1.**
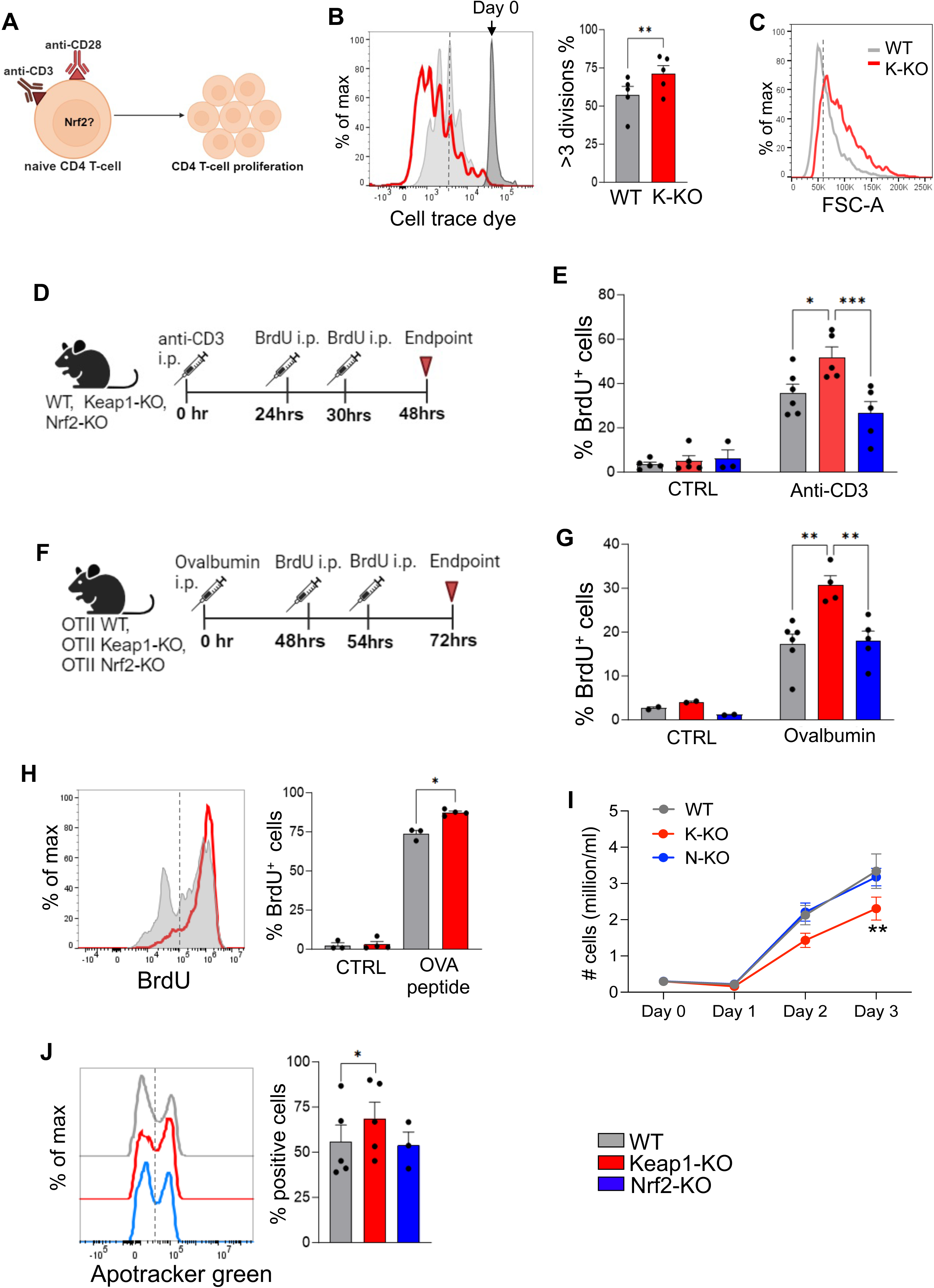
High Nrf2 promotes activation-driven proliferation of CD4^+^T-cells. (A) An illustration depicting the *in vitro* activation assay of naïve CD4^+^T-cells using anti-CD3 and anti-CD28 antibodies, resulting in T-cell expansion. (B) Naïve CD4^+^T-cells isolated from Keap1-KO and wild-type (WT) littermate mice were labeled with Cell Trace dye and activated *in vitro* for 96 h. A representative histogram overlay shows the dye dilution, and the graph shows the summary of proliferation (n=4-5). (C) Representative histogram overlay comparing the cell size (FSC, forward scatter) of WT and Keap1-KO CD4^+^T-cells 96h post-activation. (D) Schematic diagram of *in vivo* BrdU incorporation assay after injecting anti-CD3 antibodies into the indicated mice. (E) Summary graphs of percentages of BrdU^+^CD4^+^T-cells within mesenteric lymph nodes of WT, Keap1-KO, and Nrf2-KO mice treated as in (D) (n=3-6). (F) Schematic diagram of *in vivo* BrdU incorporation assay in ova-TCR transgenic (OTII)-WT, OTII-Keap1-KO, and OTII-Nrf2-KO mice after antigen challenge with ovalbumin. (G) Percentages of splenic BrdU^+^CD4^+^T-cells in indicated OTII mice treated as in (F) (n=4-6). (H) Total splenocytes from OTII-WT and OTII-Keap1-KO mice were cultured *in vitro* with ova peptide in the presence of BrdU. Histogram overlay and graph shows percentages of BrdU^+^CD4^+^Vβ5.1^+^T-cells (n=3-4). (I) Daily cell counts of *in vitro* activated WT, Keap1-KO, and Nrf2-KO CD4^+^T-cells were measured (n=7-11). (J) Representative histogram overlay and summary data of Apotracker green staining in CD4^+^T-cells 72h post-activation *in vitro* (n=3 or 5). All data are shown as mean ± SEM from the indicated number of sets of mice. *p<0.05, **p<0.01, ****p<0.0001. K-KO: Keap1-KO, N-KO: Nrf2-KO

### High Nrf2 enhances CD4^+^T-cell activation and TCR signaling

We next probed if higher Nrf2 activity in activated CD4^+^ T-cells also modulates their activation levels. Cell surface expression of activation markers and intracellular TCR-signaling events were compared after *in vitro* activation of WT, Keap1-KO, and Nrf2-KO CD4^+^T-cells. The expression of early activation markers CD44 and CD25 (IL-2Ra) was elevated in Keap1-KO CD4^+^T-cells (Figures 2A and 2B) while Nrf2-KO CD4^+^T-cells showed a modest decrease compared to WT cells (Figures 2A and 2B). Consistent with *in vitro* findings, OTII-Keap1-KO CD4^+^T-cells displayed increased expression of CD25 and CD44 than OTII-WT and OTII-Nrf2-KO CD4^+^T-cells upon ovalbumin challenge *in vivo* (Figures 2C and 2D) suggesting enhanced activation of CD4^+^T-cells with high Nrf2 expression.

**Figure 2.**
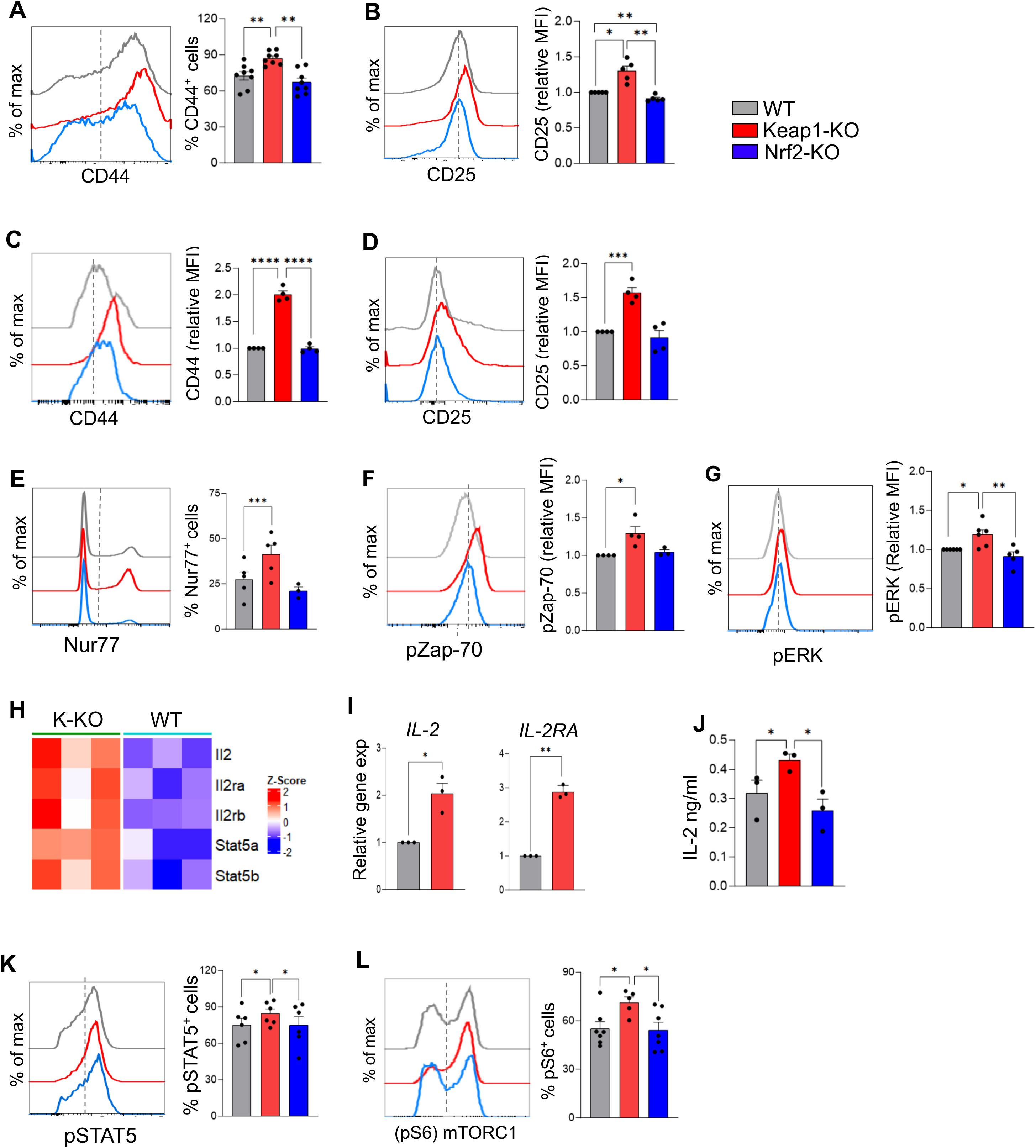
High Nrf2 enhances CD4^+^T-cell activation and TCR signaling. (A, B) The expression of (A) CD44 (n=8) and (B) CD25 (n=5) was measured in naïve CD4^+^T-cells from WT, Keap1-KO, and Nrf2-KO mice activated *in vitro* as in Figure 1A. (C, D) OTII-WT, OTII-Keap1-KO, and OTII-Nrf2-KO mice were challenged with ovalbumin *in vivo*. The expression of (C) CD44, and (D) CD25 in CD4^+^ Vβ5.1^+^ (OTII) cells was measured (n=4-5). (E-G) The expression of Nur77 (E) after 4h, and phosphorylated Zap-70 (F) and ERK (G) after 24h was compared between WT, Keap1-KO, and Nrf2-KO CD4^+^T-cells activated *in vitro* (n=3-6). (H) Heatmap of RNAseq analysis comparing the gene expression levels in Keap1-KO vs WT naïve CD4^+^T-cells 24h post-activation (n=3). (I) Relative gene expression levels of *IL-2* and *IL-2ra* in WT and Keap1-KO CD4^+^T-cells 24h post-activation measured by qRT-PCR. Data are representative of three independent experiments. (J) Secreted IL-2 protein levels from the culture supernatants of WT, Keap1-KO, and Nrf2-KO CD4^+^T-cells 48h post-activation were measured by ELISA (n=3). (K, L) Phosphorylation levels of STAT5 (K) and S6 proteins in WT, Keap1-KO, and Nrf2-KO CD4^+^T-cells 24h post-activation are shown (n=5-7). Histogram overlays show representative data and graphs show mean ± SEM. *p<0.05, **p<0.01, ****p<0.0001.

Activation events are a consequence of intracellular signaling cascades^35,36^ triggered by antigen-bound TCR complexes^37^. Our GO enrichment analysis identified upregulated expression of gene sets belonging to protein serine/threonine kinase and kinase regulator activity in activated Keap1-KO CD4^+^T-cells (Figures S2A and S2B). Therefore, we examined the expression of the orphan nuclear receptor Nur77 and the phosphorylation status of protein tyrosine kinase Zap-70, MAP Kinase pathway protein ERK and NF-kB, all of which are key components of the TCR signaling cascade^38,39,40^. The expression of Nur77 and pZAP-70 was upregulated in Keap1-KO CD4^+^T-cells while the WT and Nrf2-KO CD4^+^T-cells were similar (Figures 2E and 2F). Distal TCR signaling proteins pERK and pNF-kB were also heightened in Keap1-KO CD4^+^T-cells (Figures 2G and S2C). Together, these data indicate that high Nrf2 levels potentiate a stronger TCR signal in activated CD4^+^T-cells.

TCR signaling and costimulatory signals induce upregulation of IL-2R and interleukin-2 (IL-2) secretion, which are indispensable for T-cell expansion^41,42^. The RNAseq data revealed upregulated expression of *IL-2* and IL-2 receptor genes *Il2ra* and *Il2rb* in addition to other genes associated with IL-2 signaling like *Stat5a and Stat5b* in Keap1-KO CD4^+^T-cells (Figure 2H). Increased expression of *Il-2* and *Il-2ra (CD25)* were validated by quantitative RT-PCR (Figure 2I). Consistently, secreted IL-2 protein levels were also higher in the Keap1-KO CD4^+^T-cell culture supernatants than in WT and Nrf2-KO cultures (Figure 2J). Further, Keap1-KO CD4^+^T-cells also exhibited upregulated activity of pSTAT5, a downstream target of IL-2 signaling^43^. We next measured the levels of mammalian/mechanistic target of rapamycin complex1 (mTORC1), a metabolic sensor that integrates environmental cues for T cell activation, expansion, and function^44^ and found it increased in activated Keap1-KO CD4^+^T-cells (Figure 2K). (Figure 2L). Taken together, these findings indicate that high Nrf2 activity in activated CD4^+^T-cells enhances the overall activation by upregulating TCR-and IL-2 signaling and may lead to their superior expansion by mTORC1-driven metabolic reprogramming.

### Nrf2 suppresses glucose metabolism in activated CD4^+^ T-cells

Because glycolysis fuels activation-driven expansion^4,6,45^, we reasoned that the heightened proliferation rate of Keap1-KO CD4^+^T-cells is supported by increased glycolysis. We and others have previously shown high Nrf2 activity in Keap1-KO T-cells^25,32^. To validate that the metabolic defects observed in Keap1-KO CD4^+^T-cells are mediated by the increased Nrf2, and not due to the deficiency of Keap1, we included in our study, mice depleted of both Keap1 and Nrf2 in T-cells (Keap1^fl/fl^Nrf2^fl/fl-CD4Cre^, called double-KO; D-KO) as we reported previously^25^. As hypothesized the proliferative capacity of activated Double-KO CD4^+^T-cells is restored to WT levels (Figure S3A). To assess if high Nrf2 in CD4^+^T-cells upregulates glycolysis, we assessed the levels of lactate, a byproduct of glycolysis^46,47^. Contrary to our hypothesis, the levels of intracellular and extracellular lactate (in supernatants) were lower in activated Keap1-KO than in WT CD4^+^T-cells (Figure 3A) but were restored to WT levels in double-KO CD4^+^T-cells. Pyruvate, which is converted to lactate was also decreased in the Keap1-KO CD4^+^T-cells (Figure S3B). We then performed a glycolytic stress test using activated WT, Keap1-KO, and double-KO CD4^+^T-cells and found a lower extracellular acidification rate (ECAR) in Keap1-KO CD4^+^T-cells compared to the WT cells, which was restored partially in the double-KO CD4^+^T-cells (Figures 3B and 3C). Keap1-KO CD4^+^T-cells also had decreased glycolytic capacity (Figures 3D) and glycolytic ATP (Figure 3E). Consistently, ova-specific stimulation of OTII-Keap1-KO CD4^+^T-cells *in vivo* also showed lower ECAR and glycolytic capacity than OTII-WT CD4^+^T-cells (Figures 3F-H). These data suggest that activated Keap1-KO CD4^+^T-cells have a low glycolysis rate and might not rely on glucose-fueled glycolysis for their hyper-proliferation. To prove this, we performed *in vitro* activation of WT and Keap1-KO CD4^+^T-cells in glucose-depletion conditions and measured proliferation. As seen earlier (Figure 1B), Keap1-KO CD4^+^T-cells proliferated better than WT cells in normal media containing glucose (Figure 3I, upper panel). In the absence of glucose however, WT CD4^+^T-cells proliferated poorly as expected, but the Keap1-KO CD4^+^T-cells continued to proliferate at an increased rate (Figure 3I, lower panel). Taken together, these findings suggest that high Nrf2 in CD4^+^ T-cells suppresses glycolysis and supports their proliferation in a glucose-independent manner.

**Figure 3.**
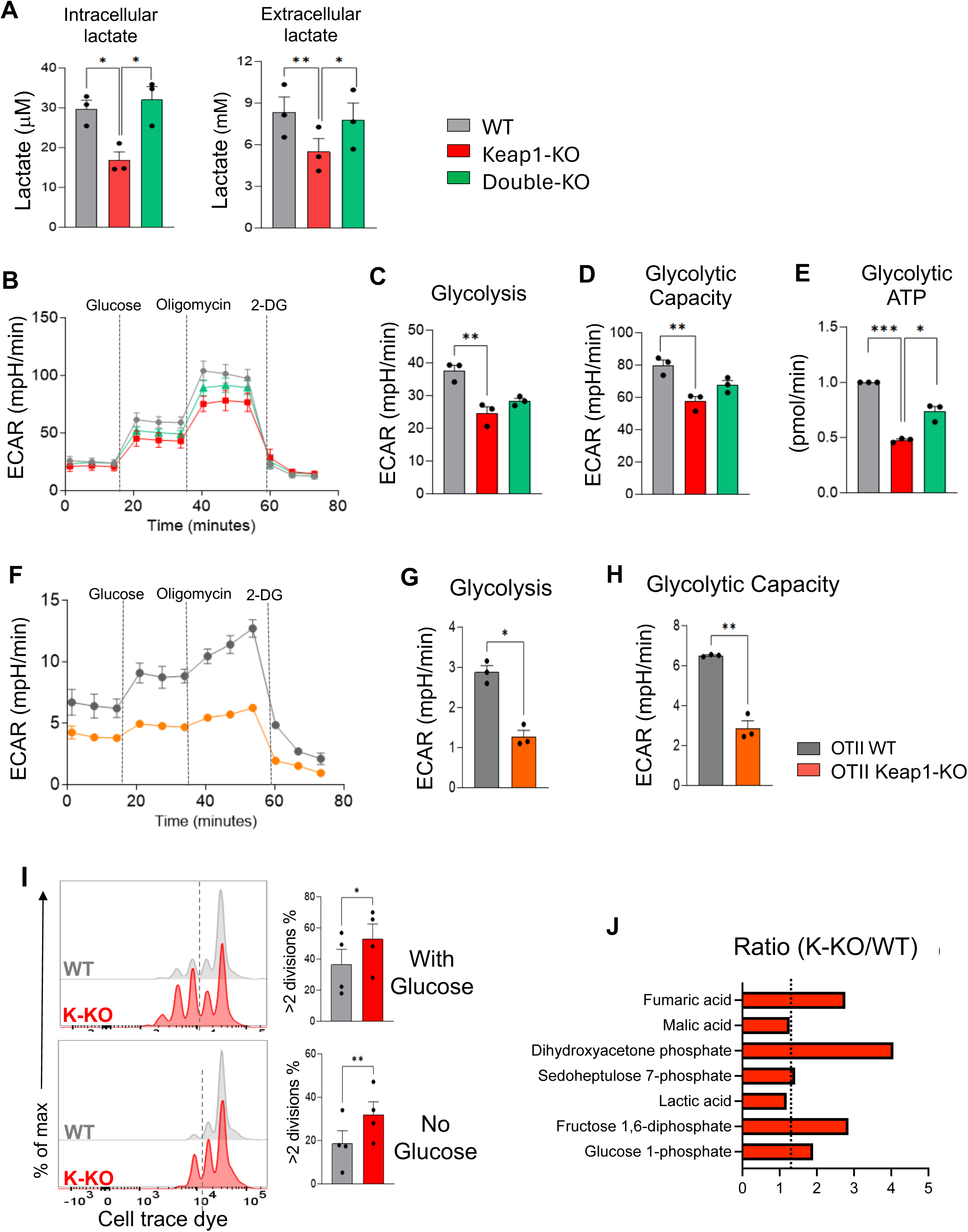
Nrf2 suppresses glucose metabolism in activated CD4^+^T-cells. (A) Levels of intracellular and extracellular lactate from WT, Keap1-KO, and Double-KO CD4^+^T-cells were measured 48h post-activation *in vitro* (n=3). (B-E) Naïve CD4^+^T-cells from WT, Keap1-KO, and Double-KO mice were activated *in vitro* for 48h and subjected to a glycolytic stress test. The representative graphs show ECAR over time (B), glycolysis (C), glycolytic capacity (D), and glycolytic ATP (E) (n=3). (F-H) A glycolytic stress test was performed using OTII-WT and OTII-Keap1-KO CD4^+^T-cells isolated from mice 72h after the ova challenge. ECAR over time (F), glycolysis (G), and glycolytic capacity (H) are shown. Data is representative of two independent experiments. (I) Representative histogram overlay and summary of the proliferation of WT and Keap1-KO CD4^+^T-cells activated *in vitro* in media with 11mM glucose (*upper panel*) or without glucose (*lower* panel) (n=4). (J) The ratio of indicated metabolites in Keap1-KO over WT CD4^+^T-cells activated *in vitro* for 72 h followed by CE-MS analysis (n=2). Data are shown as mean ± SEM. *p<0.05, **p<0.01, ****p<0.0001. K-KO: Keap1-KO

To understand Nrf2-driven nutrient metabolism in CD4^+^T-cells, we performed metabolomics to assess total metabolite concentrations. The ratio of metabolites in Keap1-KO over WT CD4^+^T-cells is shown in Figure 3J. The early glycolytic six-carbon metabolites (Glucose 1-phosphate and Fructose 1,6-diphosphate) were higher in K-KO CD4^+^T-cells, but the three-carbon lactate production was reduced, indicating that glucose oxidation shifted away from lactate in these cells. The glucose uptake between the two cell types was comparable (Figure S3B) suggesting that the glucose likely feeds into alternative metabolic pathways. Indeed, the levels of Dihydroxyacetone Phosphate (DHAP) were higher in K-KO CD4^+^T-cells suggesting an increase in glucose-fueled triglyceride synthesis (Figure 3J)^48^. In addition, the TCA intermediate Fumaric acid was higher than WT in K-KO CD4^+^T-cells (Figure 3J) despite lower pyruvate (Figure S3B). In line with the metabolite data, Keap1-KO CD4^+^T-cells showed an upregulation of the genes involved in the initial steps of glucose metabolism like hexokinase1 (*HK1*), phosphofructokinase, muscle (*PFKM*), aldolase, and fructose-bisphosphate C (*ALDOC*), in addition to the enzymes involved in PPP like glucose-6-phosphate dehydrogenase (*G6pd*) and transaldolase 1 (*TALDO1*)^49^ (Figures S2A and S3D). Overall, these data suggest low glycolysis but a carbon flux into TCA metabolism and ATP production from a non-glucose source, probably glutamine in activated CD4^+^T-cells with high Nrf2.

### Nrf2 promotes glutamine metabolism to support CD4^+^ T-cell proliferation

CD4^+^T-cells upregulate the expression of Glutamine (Gln) transporters and rely on Gln for activation and IL-2 expression^9,50,51,52^. Higher amounts of intracellular Gln were noted in activated Keap1-KO compared to the WT and double-KO CD4^+^T-cells (Figure 4A). The fact that Nrf2 is a positive regulator of Gln metabolism in cancer cells^53,54^ prompted us to hypothesize that activated Keap1-KO CD4^+^ T-cells upregulate Gln metabolism to support their proliferation. Glutaminolysis involves the conversion of glutamine to glutamate and then to α-ketoglutarate (α-KG), which may then enter the TCA cycle to generate ATP^50,55,56^. As hypothesized, we noted higher levels of glutamate and α-KG in Keap1-KO CD4 ^+^T-cells, which were restored to WT levels in the double-KO CD4^+^T-cells (Figure 4B and 4C). To further examine the Gln utilization pathways in high-Nrf2 CD4^+^ T-cells, we performed carbon-tracing experiments using carbon13 isotope-labeled Gln ([U-^13^C]-Gln) followed by CEMS-based metabolomics as done before^57^. We noticed more C13 isotope incorporation (from C13-labeled Gln) than C12 (from non-labeled glucose) into the total metabolites in the Keap1-KO CD4^+^T-cells, which was not the case with WT cells indicating higher Gln metabolization by these cells (Figure 4D). Citrate is an intermediate of the TCA cycle which is fed either by Acetyl-CoA via glycolysis and glucose oxidation or by the oxaloacetate via α-KG and TCA cycle^58^. Gln-tracing also revealed a higher abundance of C13-derived citrate in activated Keap1-KO CD4^+^T-cells than in WT cells (Figure 4E) as measured by the levels of M4 isotopomers which indicate oxidative TCA metabolism from glutamine. Thus, unlike the WT CD4^+^T-cells, Nrf2 drives an increase in Gln-mediated TCA flux in Keap1-KO CD4^+^T-cells.

**Figure 4.**
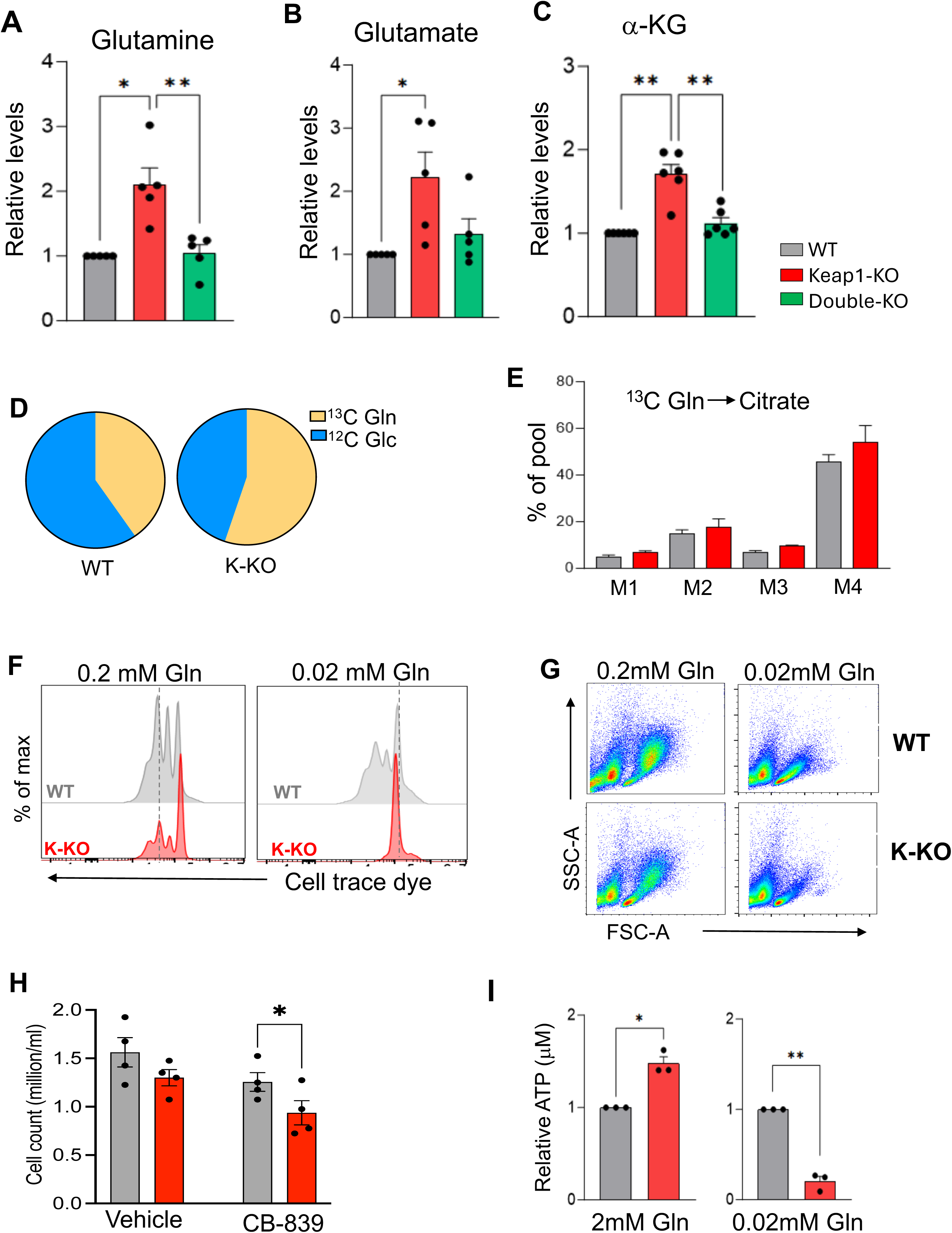
Nrf2 promotes glutamine metabolism to support CD4^+^T-cell proliferation. (A-C) The Levels of glutamine (A), glutamate (B), and a-ketoglutarate (C) from WT, Keap1-KO, and Double-KO naïve CD4^+^T-cells were measured 48h after *in vitro* stimulation (n=5-6) (D) The percentage contributions of ^13^C-labeled glutamine (M1+) and ^12^C Glucose (M0) to the total metabolites isolated from activated WT and Keap1-KO CD4^+^T-cells are shown. (n=2). (E) Graph shows the % pool of Citrate from ^13^C-labeled glutamine in WT, Keap1-KO CD4^+^T-cells 72 h post-activation (n=2). (F-G) Representative plots show the proliferation (A) and the cell size (G) of WT and Keap1-KO CD4^+^T-cells activated *in vitro* in media containing limiting glutamine (0.2mM or 0.02mM) and glucose (n=4). (H) WT and Keap1-KO CD4^+^T-cells were treated with Vehicle or 250nM CB-839 for xx h during *in vitro* activation. Summary graphs compare the cell counts (n=4). (I) Intracellular ATP levels were compared between WT and Keap1-KO CD4^+^T-cells activated in media with glucose and 2mM or 0.02mM glutamine (n=3). Data are shown as mean ± SEM. *p<0.05, **p<0.01, ****p<0.0001. K-KO: Keap1-KO, Gln: glutamine, Glc: glucose

Next, we asked if the activated Keap1-KO CD4^+^T-cells rely on Gln as their primary carbon source for increased proliferation by analyzing the activation status and rate of proliferation in Gln-limiting conditions. While limiting Gln reduced the overall proliferation of both cell types, Keap1-KO CD4^+^T-cells lost their ability to proliferate at an increased rate in low (0.2mM and 0.02mM) glutamine conditions (Figures 4F and 4G). They depicted a complete ablation of proliferation at 0.02 mM Gln despite having sufficient glucose in the media (Figure S4A and 4F). The expression of activation markers, CD69, CD25, and CD44 by Keap1-KO CD4^+^T-cells was also severely dampened at low Gln concentrations (Figure S4B). Then, we probed if inhibiting glutaminolysis abrogates the superior proliferation capacity of Keap1-KO CD4^+^T-cells using the glutaminase inhibitor CB-839, which inhibits the first step of glutaminolysis^59^. CB-839 has a more robust effect on the proliferation and cell numbers of activated Keap1-KO CD4^+^T-cells than on WT cells (Figures 4H and S4C). Lastly, to probe if Nrf2-high CD4 T-cells depend on Gln as their energy source, we measured the intracellular ATP levels of WT and Keap1-KO CD4^+^T-cells activated in Gln-sufficient (2mM) and Gln-low (0.02mM) conditions and found significantly low ATP in Keap1-KO CD4^+^T-cells activated in Gln-low conditions implying that they derive their ATP from Gln, and not from glucose (Figure 4I). In summary, our results suggest that high Nrf2 in activated CD4^+^T-cells favors glutamine metabolism to support increased CD4^+^T-cell proliferation.

## DISCUSSION

A few previous studies mostly using Nrf2-activating drugs show that Nrf2 regulates T-cell activation, primarily through its involvement in redox homeostasis and antioxidant defense mechanisms^26,28,30,60^ but the underlying mechanisms or the impact on T-cell expansion is unknown. Here, using mice with T-cell-specific deletion of Keap1 or Nrf2, we systematically studied the effects of high and low Nrf2, respectively^32^ on T-cells. We report that Keap1-KO CD4^+^T-cells with constitutively high Nrf2 activity exhibit an increase in activation-driven proliferation *in vitro* and *in vivo* while Nrf2-KO CD4^+^T-cells exhibit a modest decrease. Keap1-KO mice with high Nrf2 in T-cells did not show a difference in T-cell development or cell frequencies as reported by us and others^25,32^indicating that Nrf2 specifically drives their activation-driven expansion., Nrf2 gene expression and protein levels were found to be upregulated in human T-cells during the initial hours of stimulation^31^. We similarly noted increased Nrf2 protein expression during early activation in CD4^+^T-cells from a C57B/L6 mouse, but interestingly, the expression decreased at later stages when the cells were in the log phase of robust proliferation. Thus, while Nrf2 likely plays a crucial role in the initial activation of T-cells, its sustained activity may potentially impede late-stage processes, leading to its downregulation.

TCR signaling events initiated by the antigen recognition lead to T-cell activation and clonal expansion^4,35^ by upregulation of autocrine IL-2 production. The phosphorylation activity of both proximal (Zap-70) and distal kinases (ERK, S6, NF-kB) involved in the TCR signaling was increased with higher Nrf2 activity but not different with Nrf2 deletion implying a stronger TCR signaling in Keap1-KO CD4^+^T-cells in response to activation. In addition, these cells show increased IL-2 and IL-2 signaling, correlating with their hyperproliferative capacity^43,52,61^. Nrf2 may promote IL-2 expression likely by direct transcriptional activation of *Il-2* and related genes. However, the crosstalk between Nrf2 and TCR signaling is yet to be determined.

To meet the rapid energy requirement of clonal expansion, naïve T-cells switch from OXPHOS to glycolysis^45,62,63^ driven by mTORC1^44,64^. Our findings show increased proliferation together with elevated mTORC1 activity in activated Keap1-KO CD4^+^T-cells prompting us to speculate that Nrf2 probably upregulates glycolysis as a metabolic mechanism for driving the activation-driven expansion of these cells. On the contrary, Keap1-KO CD4^+^T-cells had low levels of lactate and decreased ECAR indicating lower glycolysis compared to WT cells. Intriguingly, metabolomics data depicted increased levels of early glycolytic intermediates (Glucose 1-phosphate and Fructose 1,6-diphosphate) with high Nrf2. Comparable glucose uptake between activated WT and Keap1-KO CD4^+^T-cells indicated that glucose is likely being metabolized through an alternate pathway other than glycolysis in these Nrf2-high CD4^+^T-cells. Indeed, increased levels of DHAP and sedoheptulose 7-phoshate, indicative of gluconeogenesis and pentose phosphate pathway (PPP), respectively were observed in Keap1-KO CD4^+^T-cells which likely generate the reducing equivalents like NADPH for the enhanced antioxidant functions^65^. Keap1-KO CD4^+^T-cells did not rely on glucose for their hyper-proliferative capacity suggesting that Nrf2 expression levels in activated CD4^+^T-cells impart metabolic flexibility in the utilization of glucose. While this needs further investigation, in the current study, we focused on identifying the alternate nutrient source used by the Nrf2-high CD4^+^T-cells for supporting the increased proliferation.

Glutamine is the most abundant amino acid in the blood and its deficiency impacts immune cell functions^45,50,66^ including T-cells which rely on Gln for activation and proliferation^4,9,67^. We speculated that akin to rapidly dividing cancer cells, Nrf2 likely imparts a Gln-addiction in Keap1-KO CD4^+^T-cells^53,54^. Accordingly, the levels of Gln, glutamate, and α-KG were high in these cells. α-KG influences the mTOR signaling pathway and affects the process of T-cell differentiation^68^ which might explain why mTORC1 activity is elevated in Nrf2-high CD4^+^T-cells. Moreover, α-KG can transfer carbon atoms and electrons into the TCA cycle, which is central to cellular respiration and energy production. Consistent with this, Gln-tracing experiments revealed that Gln predominantly contributes to the production of TCA intermediates citrate and fumarate in Keap1-KO CD4^+^T-cells, unlike in WT CD4^+^T-cells which derive them mostly from glucose suggesting an augmented carbon flow into the TCA cycle in CD4^+^T-cells with high Nrf2. Understandably so, limiting Gln in the culture media hampered the ATP production of Nrf2-high Keap1-KO CD4^+^T-cells much more significantly than that of the WT CD4^+^T-cells with regulated Nrf2. In addition, limiting Gln also impaired the proliferation, affected the cell size, and dampened the activation status of Keap1-KO CD4^+^T-cells significantly despite the presence of glucose in the media, indicating that Nrf2-high CD4^+^T-cells rely heavily on Gln over glucose for supporting CD4^+^T-cell expansion. Unclear at this point, understanding the transcriptional regulation of glutamine metabolism by Nrf2 in activated CD4^+^T-cells can further provide valuable insights into how Nrf2 influences the metabolic state of T-cells.

With a myriad of Nrf2-activating drugs being tested in pre-clinical and clinical trials for various diseases and inflammatory conditions^60,69–71^, it is highly significant to understand how high Nrf2 affects T-cell activation and their immunometabolism. In summary, we uncover an important role of Nrf2 in regulating the nutrient dependency and metabolism of activated CD4^+^T-cells to govern their activation and expansion thus paving the way for the utilization of metabolism as a tool for improving the existing Nrf2-based therapeutics including T-cell immunotherapies.

## ACKNOWLEDGEMENTS

We thank Dr. Cheong-Hee Chang for her guidance and generous support for this work. We thank Ms. Emily Burt and Ms. Julie Hix for diligently maintaining the mouse colonies for this study. We thank Dr. Alexander Buko (HMT America, Inc) for assistance with metabolomics data analysis and scientific editing. This work was supported by startup funds to K.P. by Kansas State University (KSU) and the University of Kansas Cancer Center (KUCC). We acknowledge the support and funds provided by NIGMS (K-INBRE, P20 GM103418) to K.P, A.P, and A.B. We also acknowledge the funds to K.P. by Kansas Institute for Precision Medicine from the NIGMS (P20 GM130423). D.D. received a graduate student cancer research award from Johnson Cancer Research Center of KSU. We acknowledge the Flow Cytometry Core at KUMC, supported, in part, by the NIH/NIGMS COBRE grant P30 GM103326 and the NIH/NCI Cancer Center grant P30 CA168524.

## AUTHOR CONTRIBUTIONS

K.P. designed and supervised the study. A.T., D.D., A.P., A.B., and B.C. performed experiments and data analysis. N.K.Y. analyzed the transcriptomics data. S.G. helped in performing Seahorse experiments and in editing the manuscript. A.T. and K.P. wrote the manuscript.

## STAR Methods

### EXPERIMENTAL MODEL AND SUBJECT DETAILS

#### Mice strains and maintenance

Keap1^fl/fl-CD4Cre^ mice (referred to as Keap1-KO; K-KO) generated as described previously^25,32^ were kindly provided by Dr. Hamid Rabb (Johns Hopkins University). Nrf2^fl/fl-CD4Cre^ (Nrf2-KO; N-KO) mice were generated by crossing Nrf2^fl/fl^ mice^72^ (provided by Dr. Sekhar P. Reddy, University of Illinois) with CD4-Cre mice (The Jackson Laboratory). The Keap1 and Nrf2 double knock-out (Keap1^fl/fl^Nrf2^fl/fl-CD4Cre^; D-KO) mice were generated by crossing Keap1^fl/flCD4Cre^ mice with Nrf2^fl/flCD4Cre^ mice. The OT-II-Keap1^fl/fl-CD4Cre^ and OT-II-Nrf2^fl/fl-CD4Cre^ mice were generated by crossing OT-II mice (The Jackson Laboratory) with Keap1^fl/flCD4Cre^ mice with Nrf2^fl/flCD4Cre^ mice, respectively. All the animals were bred and maintained at the animal care facility at the Kansas State University (KSU) and the University of Kansas Medical Center (KUMC) under specific pathogen-free conditions. For all experiments, age-matched (6-12 weeks of age) from Keap1^fl/fl^ or Nrf2^fl/fl^ littermate mice without the Cre-transgene were used as wildtype (WT) controls. All the procedures for the handling of animals were approved by the Institutional Animal Care and Use Committee at KSU (PI: Pyaram, Protocol # 4424), and KUMC (PI: Pyaram, Protocol # 23-02-298).

#### *In vivo* T-cell activation

Eight-week-old Keap1-KO, Nrf2-KO and WT littermate mice were injected with 50mg of anti-CD3 antibody (Biolegend) in 0.25 ml of sterile PBS intraperitoneally (i.p.). Control mice were injected with 0.25ml PBS. After 24 hours, mice were injected twice with 0.2ml of 2.5mg/ml of BrdU (BD Biosciences) i.p. as mentioned in BrdU incorporation assay.

#### Ova antigen challenge

Eight-week-old OTII-Keap1-KO, OTII-Nrf2-KO, and OTII-WT littermate mice were injected intraperitoneally with 100mg of Ovalbumin (Sigma) in 100ml sterile 1x PBS volume mixed with 100ml of complete Freund adjuvant (Sigma) Control mice were injected with 0.2ml PBS. After 48 hours, ovalbumin and PBS-treated mice are intraperitoneally injected with 0.2ml of 2.5mg/ml of BrdU (BD Biosciences) as mentioned in BrdU incorporation assay.

#### *In Vivo* BrdU Incorporation

Eight-weeks old Keap1-KO, Nrf2-KO and WT littermate mice injected with anti-CD3 antibodies or challenged with Ovalbumin were intraperitoneally injected with 0.5 mg BrdU (BD Biosciences) in 0.2ml of PBS twice every six hours. The control mice received PBS. Eighteen hours after final dose, mice were euthanized and single-cell suspensions were made from the spleens, and the mesenteric lymph nodes. Cells were stained for surface markers, followed by anti-BrdU antibody to assess BrdU incorporation using BrdU flow kit (BD Biosciences), according to the manufacturer’s protocol.

### METHOD DETAILS

#### Cell preparation and purification

On the day of culture, the mice were euthanized, and the spleens were harvested. The homogenized spleen cells were then subjected to RBC lysis buffer, cells were washed, and then resuspended in 1x PBS supplemented with 1% FBS (FACS buffer). The naïve CD4 T-cells were purified using the EasySep™ Mouse Naïve CD4+ T Cell Isolation Kit (STEMCELL Technologies) as per the manufacturer’s instructions. The total CD4 T-cells were isolated using EasySep™ Mouse CD4 Positive Selection Kit II (STEMCELL Technologies). Wherever indicated, freshly prepared whole splenocytes were also used.

#### T-cell activation and *in vitro* culture

Freshly isolated and purified CD4 T cells (1 × 10^6^ cells/ml) were stimulated with plate-bound anti mouse-CD3 antibody (5 μg/ml; Invitrogen) and soluble anti mouse-CD28 Ab (1 μg/ml; Invitrogen) in RPMI-1640 (Gibco) supplemented with 10% FBS (Gibco), and 1x Penicillin/Streptomycin/Glutamine solution (Cytiva) and b-mercaptoethanol (Gibco). The cells were then cultured for the indicated amounts of time in a 37°C humidified incubator with CO_2_ until analysis.

To measure activation-driven cell proliferation, naive or total CD4 T-cells were labeled with 5mM CellTrace^TM^ Violet (CTV; Invitrogen) in 1XPBS containing 0.1% BSA for 20 min at 37C and activated with in vitro using anti-CD3 and anti-CD28 antibodies. Cells were analyzed by flow cytometry on days 2-4 to assess dilution of CTV.

For the nutrient-limiting culture experiments, RPMI 1640 without glucose or glutamine (Elabscience). For cultures without glucose, media was supplemented with 2mM L-glutamine (Gibco), 10% dialyzed FBS (Millipore Sigma) and 1x Penicillin/Streptomycin. For cultures with limiting glutamine, media was supplemented with 11mM glucose (Gibco), 0.2 mM or 0.02 mM L-glutamine, 10% dialyzed FBS (Millipore Sigma), and 1x Penicillin/Streptomycin. Media containing both 11mM glucose and 2mM L-glutamine was used as a positive control. Freshly isolated and purified CD4 T cells (1 × 10^6^ cells/ml) from Keap1-KO and WT mice were labelled with cell trace dye and stimulated with anti-CD3 and anti-CD28 antibodies in the indicated nutrient-limiting or control media for 3-4 days to assess proliferation and activation markers by flow cytometry as well as ATP levels.

#### *In Vitro* BrdU incorporation assay

Single-cell suspension of spleens was prepared from six-week-old OTII-Keap1-KO and OTII-WT littermate mice. 3 × 10^5^ splenocytes per well were seeded in 96-well round bottom plate and 5ug OVA 323-339 peptide/ml (InvivoGen) was added to the experimental groups only.

Both control and experimental were cultured with 10ul/ml of 1mM BrdU in RPMI1640 making it at a final concentration of 10uM. On day 2 cells were harvested and stained for BrdU using BrdU flow kit (BD Biosciences), according to the manufacturer’s protocol.

#### Flow Cytometry

For staining of surface molecules, up to one million cells per sample were resuspended in 100 μl of FACS buffer (1% FBS in 1x PBS) and stained with different fluorophore-conjugated anti-mouse antibodies as provided in the key resource table including but not limited to TCR-β-Pacific Blue or TCR-β-FITC, CD4-APC-Cy7, CD44-FITC or CD44-APC, CD25-APC or CD25-PE-Dazzle, CD69-Pe-Cy7, Vβ5.1-FITC or Vβ5.1-Pe-Cy7. For assessing the expression of nuclear proteins, the cells were fixed using Foxp3/Transcription buffer staining set (Invitrogen) and stained with Nur77-PE or Nrf2-PE. For the measurement of phosphorylation events, cells were fixed using Cytofix/Cytoperm Plus (BD Biosciences), and permeabilized with ice-cold methanol before staining with pS6-PeCy7, pERK-FITC, pZap70-PE and pStat5-PE antibodies. The data were acquired on a BD LSRFortessa™ X-20 or Cytek™ Aurora flow cytometers and analyzed using FlowJo (software version 10.8.2; Becton, Dickinson & Company).

#### Glucose (2-NBDG) uptake assay

To measure the uptake of glucose, CD4 T-cells from the invitro activation cultures were harvested on day 2 and incubated with 20 μM 2-NBDG [2-(*N*-(7-Nitrobenz-2-oxa-1,3-diazol-4-yl) Amino)-2-Deoxyglucose; Cayman Chemicals) in Glucose-free RPMI media for 1 hour at 37°C. The cells were then washed once with 1x FACS buffer and then stained with appropriate surface antibodies before flow cytometry analysis.

#### Immunoblotting

Splenic CD4 T-cells from C57BL/6 mice were activated *in vitro* with anti-CD3 and anti-CD28 antibodies. Cells were harvested on days 0-3, and lysates were prepared from 1 million cells using RIPA buffer (Thermo Scientific™) supplemented with protease inhibitor (Roche). Lysates were subjected to gel electrophoresis under reducing conditions (BIO-RAD) and transferred to the PVDF membrane. After blocking with 5% skimmed milk, the membrane was incubated with the following primary antibodies-anti-mouse Nrf2 (Santa Cruz Biotechnology), anti-rabbit Keap1(D6B12; Cell Signaling Technology). Beta-actin was used as a loading control (Sigma). For secondary antibody, goat anti-mouse HRP (Thermo Scientific™) was used. Signals were detected using SuperSignal West Pico chemiluminescent substrate kit (Thermo Scientific™), and iBright imaging system. The data was analyzed using ImageJ software (Rasband, W.S., ImageJ, U.S. National Institute of Health, Maryland, USA).

#### Glycolytic stress test

Naïve CD4 T-cells isolated from WT, K-KO, and D-KO mice were activated *in vitro* using anti-CD3 and anti-CD28 antibodies in RPMI media for 72 hours as described above. Extracellular Acidification Rate (ECAR) was measured using Seahorse XF96 extracellular flux analyzer (Agilent Technology, Santa Clara, CA). Briefly, cultured CD4+ T cells were washed twice, resuspended in XF Base Media (Agilent Technologies) supplemented with 2 mM L-glutamine (Agilent Technologies), 1mM Sodium Pyruvate (Agilent Technologies) and 5 x 10^5^ cells/well were plated on extracellular flux assay plates (Agilent Technologies). Live ECAR was measured with sequential addition of 10 mM D-glucose (Agilent Technologies), 2 µM oligomycin (Sigma), and 100 µM 2-deoxyglucose (Sigma). Data were analyzed by glycolysis stress test report generator using Seahorse Wave Desktop 2.6.1 software.

#### Inhibitor treatment

To test the effect of inhibiting the glutamine metabolism, CD4 T-cells were activated *in vitro* with anti-mouse-CD3 and anti-mouse-CD28 antibodies (as described earlier) in the presence of 250 nM CB-839 (Cayman Chemicals), or DMSO in RPMI-1640 media as vehicle. The cells were cultured for the indicated amount of time and assessed for cell proliferation and activation markers using antibodies listed for flow cytometry.

#### RNA isolation and RNA-sequencing analysis

Naïve CD4 T-cells from WT and K-KO mice were activated *in vitro* (as described earlier). After 24 h, total RNA was extracted from activated CD4 T-cells using RNeasy Plus Micro kit (Qiagen) and processed at the Genomics Core Facility at the University of Kansas Medical Center. Firstly, 100ng of RNA was used to perform an Agilent TapeStation QC assay to verify the integrity of the RNA and generate a RIN value (RNA Integrity Number). Using Nova-Seq 6000 S1 200 cycle reagent kit, 25 million paired-end reads were recorded for mRNA libraries. For bioinformatics analysis, raw data underwent quality processing through fastQC^73^ and were aligned to the mouse reference genome using RSEM (RNA-Seq by Expectation-Maximization^74^, resulting in the creation of a gene count matrix. Differential expression analysis was then performed between WT and Keap1-KO groups using edgeR^75^. The differentially expressed genes (DEGs) were further analyzed for gene ontology (GO) enrichment using ClusterProfiler R-package^76^ to identify significant biological processes, molecular functions, and metabolic pathways associated with the RNASeq samples. The enrichment results were prioritized based on their significance and the top GO terms were visualized as dotplots. Subsequently, box plots were generated using ggplot2^77^ to illustrate the variability in gene expression across the groups, with mean comparison P-values annotated on the respective box plots.

#### Quantitative RT-PCR

Total RNA was extracted from the activated CD4 T cells using RNeasy Plus Micro kit (Qiagen). 50 ng of RNA was used as a template to reverse transcribe into cDNA using Oligo(dT)15 Primer (Promega) and M-MLV Reverse Transcriptase (Thermo Fisher Scientific). Applied Biosystems SYBR Select Master Mix (Thermo Fisher Scientific) and primers specific for indicated genes were used to detect the relative levels of indicated mRNAs. Beta-actin was used as an internal control for normalization and the inverse log of the ΔΔCT was then calculated to quantify the gene expression in each sample and relative gene expression is shown.

#### Measurement of metabolites and ATP

##### Lactate

Intracellular and extracellular lactate levels were measured using the L-lactate assay kit (Cayman Chemicals) as per the manufacturer’s protocol. Briefly, after 2 days of culture, 0.3 x10^5^ – 3 x 10^5^ cells were harvested and centrifuged at 10000g in 4C for 5 minutes. The supernatant was used for extracellular lactate measurement while the cell pellet was used for measuring intracellular lactate levels. Both the supernatant and the lysates were deproteinized using metaphosphoric acid and 140 μl of the reaction mixture was added to 20 μl of standard or samples in each well. The sample for the extracellular lactate assay were diluted 10-fold before use while the intracellular lactate samples were used undiluted. The reaction was initiated by adding 40 μl of enzyme mixture to each well and incubated (protected from light) at RT for 20 minutes. The fluorescence was measured at λex/em = 535/595 nm using Biotek Synergy HT Multi-Mode Microplate Reader, and the level of lactate in the sample was calculated using a standard curve generated alongside each experiment.

##### α-Ketoglutarate

A fluorometric α-ketoglutarate (α-KG) assay (Sigma) was used as per manufacturer’s instructions in a 96-well flat black bottom plate (Thermo Scientific). Briefly, 100 μl of assay buffer was added to the pellet of 100,000 cells per reaction vortexed to lyse the cells. The assay reaction was set up using 50 μl of lysate or standards and equal volume of reaction mixes and after mixing on a horizontal shaker for about 1 minute, the plate was incubated (protected from light) at 37C for 30 minutes. Fluorescence reading was taken at λex/em of 535/595 nm using Biotek Synergy HT Multi-Mode Microplate Reader, and the level of α-KG in the sample was calculated using a standard curve generated alongside each experiment.

##### Glutamine and Glutamate

The measurement of glutamine and glutamate in the cell samples was performed using a Glutamine/Glutamate-Glo™ Assay kit (Promega) according to the manufacturer’s instructions. Briefly, after *in vitro* activation for 2 days, 50,000 CD4 T-cells for each test were used per well for deproteinization using equal volume of 0.3N HCL and 450mM Tris-CL, pH 8.0. For the wells being used to determine glutamine levels, 25 μl of sample or standards were mixed with equal volume of Glutaminase Enzyme Solution (25 μl Glutaminase Buffer + 0.125 μl Glutaminase enzyme). The wells for measuring glutamate levels received 25 μl of Glutaminase Buffer only. The mixture was then incubated for 30–40 minutes at RT before adding 50 μl of glutamate detection reagent to each well. After incubation at RT for 60 minutes, the luminescence of the samples, standards, and controls was read using a luminometer. A standard curve was generated alongside each experiment to calculate the level of glutamine and glutamate in the samples.

##### Pyruvate

Intracellular pyruvate levels were measured using the Pyruvate assay kit (Cayman Chemicals). After 2 days of in vitro activation, the cell pellets from 0.3 x 10^5^ – 3 x 10^5^ CD4 T-cells were deproteinized using metaphosphoric acid. Into each well, 140 μl of the reaction mixture was added to 20 μl of standard or undiluted samples. The reaction was initiated using 20 μl of enzyme mixture and fluorescence reading was measured at λex/em = 535/595 nm after incubating (protected from light) at RT for 20 minutes. The amount of pyruvate in the sample was calculated using a standard curve generated alongside each experiment.

##### ATP

The CellTiter-Glo® Luminescent Cell Viability Assay kit (Promega) was used to measure ATP levels as per the manufacturer’s protocol. Briefly, up to 70,000 cells in 70 μl RPMI or the standards (Ambeed) in a solid white 96-well plate (Corning) were mixed with 70 μl of the CellTiter-Glo Reagent and incubated at room temperature for 10 minutes protected from light. The luminescence was measured using a luminometer and the ATP levels in the sample were calculated using a standard curve generated alongside each experiment.

#### Fluoxomics and metabolomics analysis using CE-MS

Naïve CD4 T-cells isolated from K-KO and WT littermate mice were cultured in RPMI 1640 without L-glutamine (Gibco), and with 2mM [U-^13^C5] glutamine tracer (Cambridge Isotope Laboratories), and 10% dialyzed FBS (Millipore Sigma) at a concentration of 1 million cells/ml for 72 hours. Activated cells were harvested, washed with a 5% mannitol solution, and treated with 800 μL of methanol to deactivate enzymes. Thereafter, 550 μL of Milli-Q water containing internal standards solution (from Human Metabolome Technologies) was added, and the resulting extract was centrifugated at 2300g for 5 minutes at 4C. The upper aqueous layer (800 μL) was filtered through a Milli-pore 5-kDa cutoff filter by centrifugation at 9100g for 2-3 hours at 4C to eliminate proteins. Capillary electrophoresis–mass spectrometry (CE-MS) and analysis of metabolites were performed on the collected filtrate by Human Metabolome Technologies (HMT).

#### Statistical analysis

Unless otherwise stated, the presented data was obtained from at least three biological replicates. The data for all experiments were analyzed with Prism software (Prism version 10.0.0; GraphPad Software Inc., San Diego, CA). Error bars indicate the standard deviation of the experimental replicates. Paired and unpaired Student’s *t*-test was used for comparison of two groups. 2-way ANOVA with multiple comparison using Sidak’s test was performed. We used the following convention for symbols to indicate statistical significance: ns, *P* > 0.05; *, *P* ≤ 0.05; **, *P* ≤ 0.01; ***, *P* ≤ 0.001; ****, *P* ≤ 0.0001.

#### Data and code availability

RNA sequencing data reported in the supplemental information of this publication can be accessed at the Harvard Dataverse using this link https://doi.org/10.7910/DVN/AY1YBZ. The related R-code used for the data analysis and gene ontology enrichment analysis will be provided by the lead contact upon request.

#### Key Resource Table

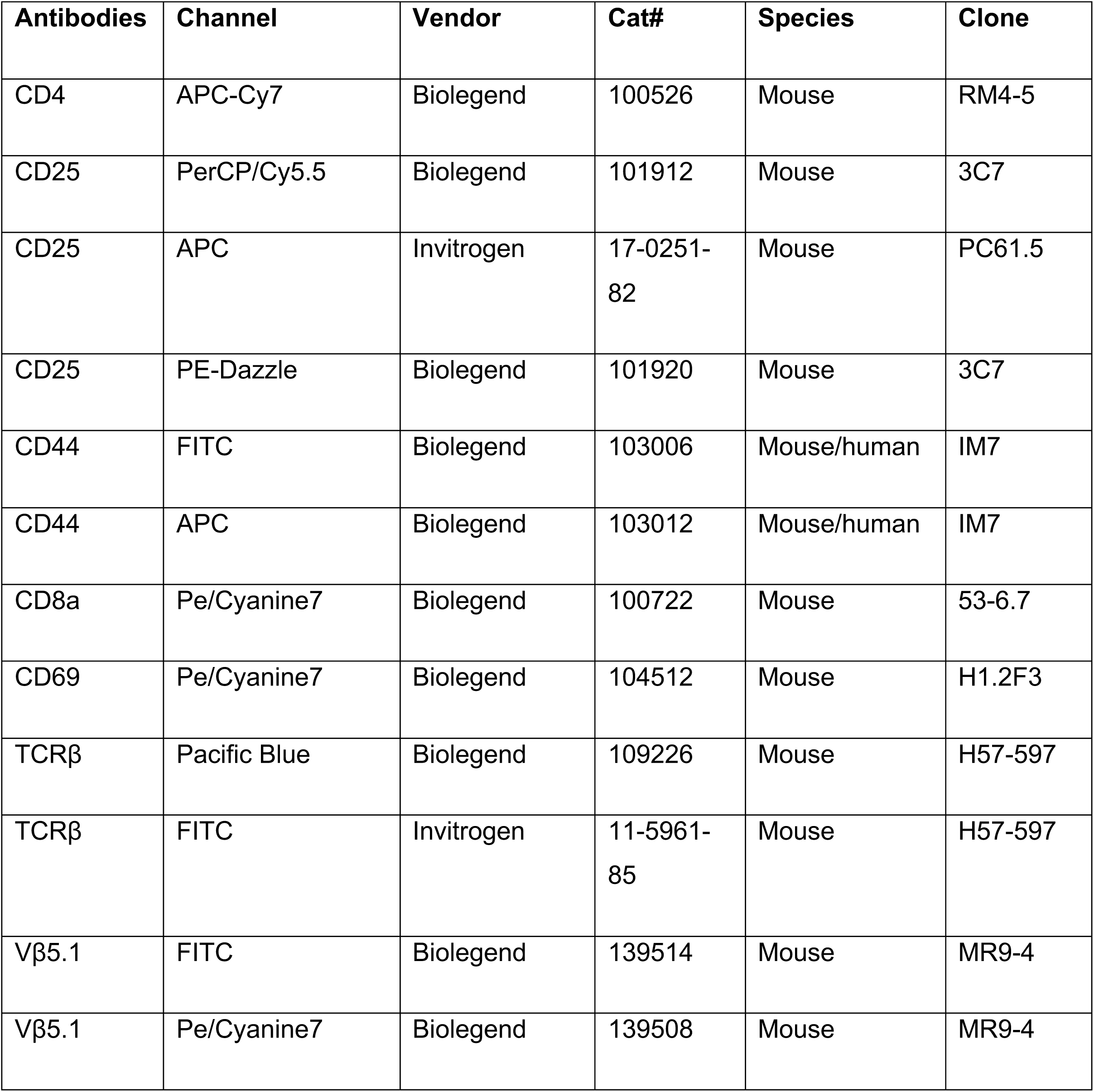
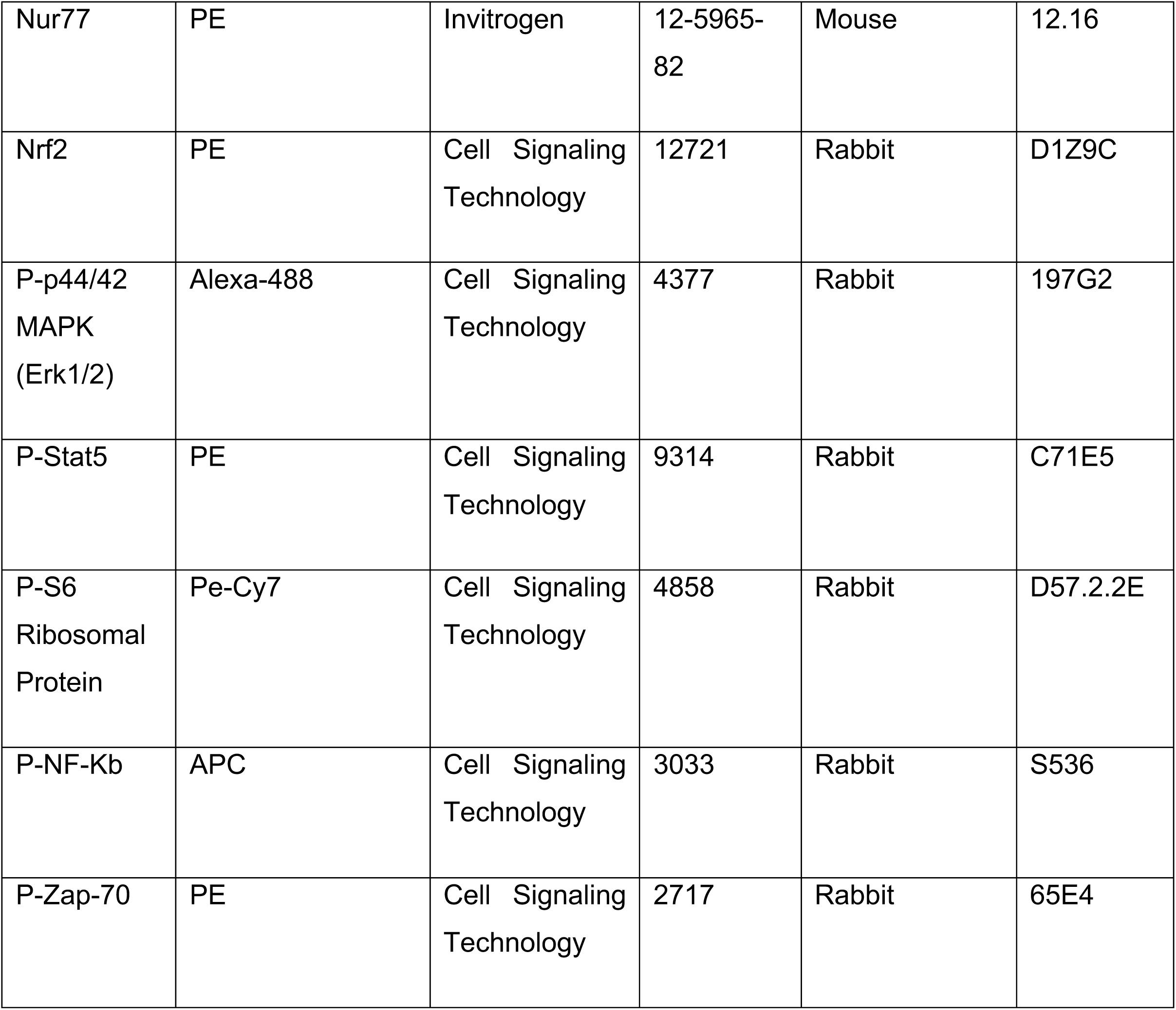
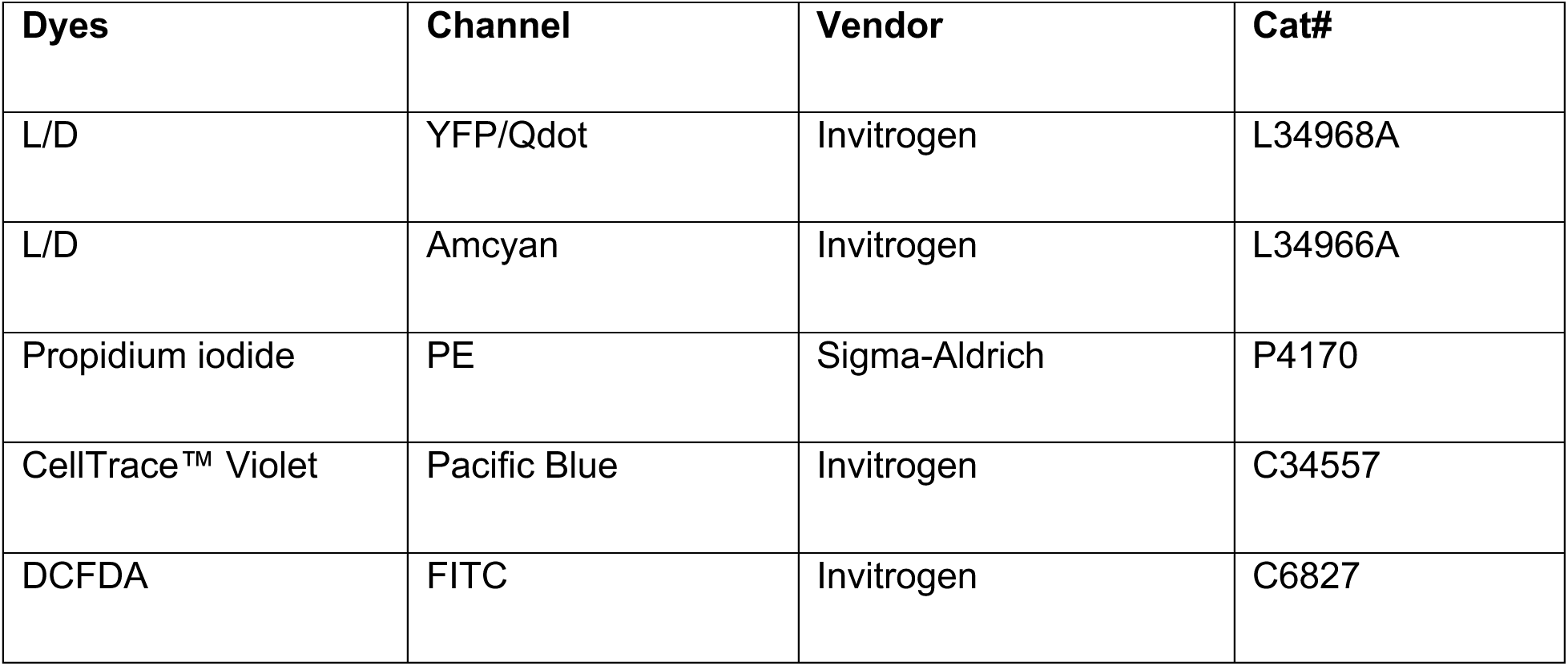
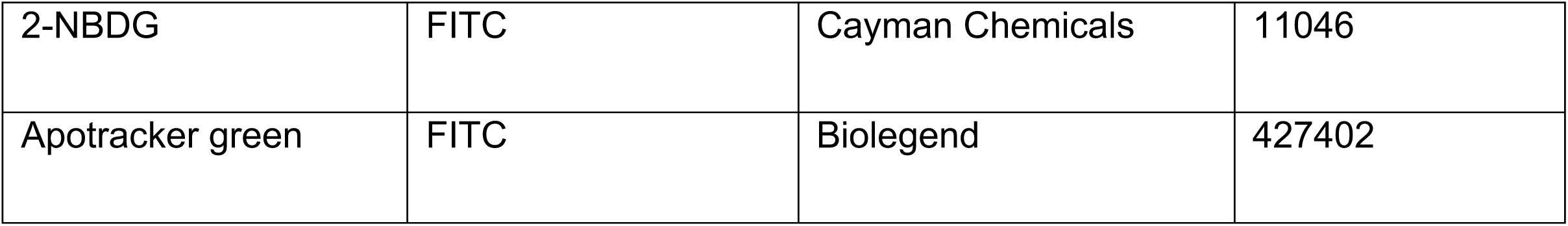

| Antibodies | Channel | Vendor | Cat# | Species | Clone |
| --- | --- | --- | --- | --- | --- |
| CD4 | APC-Cy7 | Biolegend | 100526 | Mouse | RM4-5 |
| CD25 | PerCP/Cy5.5 | Biolegend | 101912 | Mouse | 3C7 |
| CD25 | APC | Invitrogen | 17-0251-82 | Mouse | PC61.5 |
| CD25 | PE-Dazzle | Biolegend | 101920 | Mouse | 3C7 |
| CD44 | FITC | Biolegend | 103006 | Mouse/human | IM7 |
| CD44 | APC | Biolegend | 103012 | Mouse/human | IM7 |
| CD8a | Pe/Cyanine7 | Biolegend | 100722 | Mouse | 53-6.7 |
| CD69 | Pe/Cyanine7 | Biolegend | 104512 | Mouse | H1.2F3 |
| TCRβ | Pacific Blue | Biolegend | 109226 | Mouse | H57-597 |
| TCRβ | FITC | Invitrogen | 11-5961-85 | Mouse | H57-597 |
| Vβ5.1 | FITC | Biolegend | 139514 | Mouse | MR9-4 |
| Vβ5.1 | Pe/Cyanine7 | Biolegend | 139508 | Mouse | MR9-4 |
| Nur77 | PE | Invitrogen | 12-5965-82 | Mouse | 12.16 |
| Nrf2 | PE | Cell Signaling Technology | 12721 | Rabbit | D1Z9C |
| P-p44/42 MAPK (Erk1/2) | Alexa-488 | Cell Signaling Technology | 4377 | Rabbit | 197G2 |
| P-Stat5 | PE | Cell Signaling Technology | 9314 | Rabbit | C71E5 |
| P-S6 Ribosomal Protein | Pe-Cy7 | Cell Signaling Technology | 4858 | Rabbit | D57.2.2E |
| P-NF-Kb | APC | Cell Signaling Technology | 3033 | Rabbit | S536 |
| P-Zap-70 | PE | Cell Signaling Technology | 2717 | Rabbit | 65E4 |

| Dyes | Channel | Vendor | Cat# |
| --- | --- | --- | --- |
| L/D | YFP/Qdot | Invitrogen | L34968A |
| L/D | Amcyan | Invitrogen | L34966A |
| Propidium iodide | PE | Sigma-Aldrich | P4170 |
| CellTrace™ Violet | Pacific Blue | Invitrogen | C34557 |
| DCFDA | FITC | Invitrogen | C6827 |
| 2-NBDG | FITC | Cayman Chemicals | 11046 |
| Apotracker green | FITC | Biolegend | 427402 |

#### Chemicals, Peptides, and Reagents

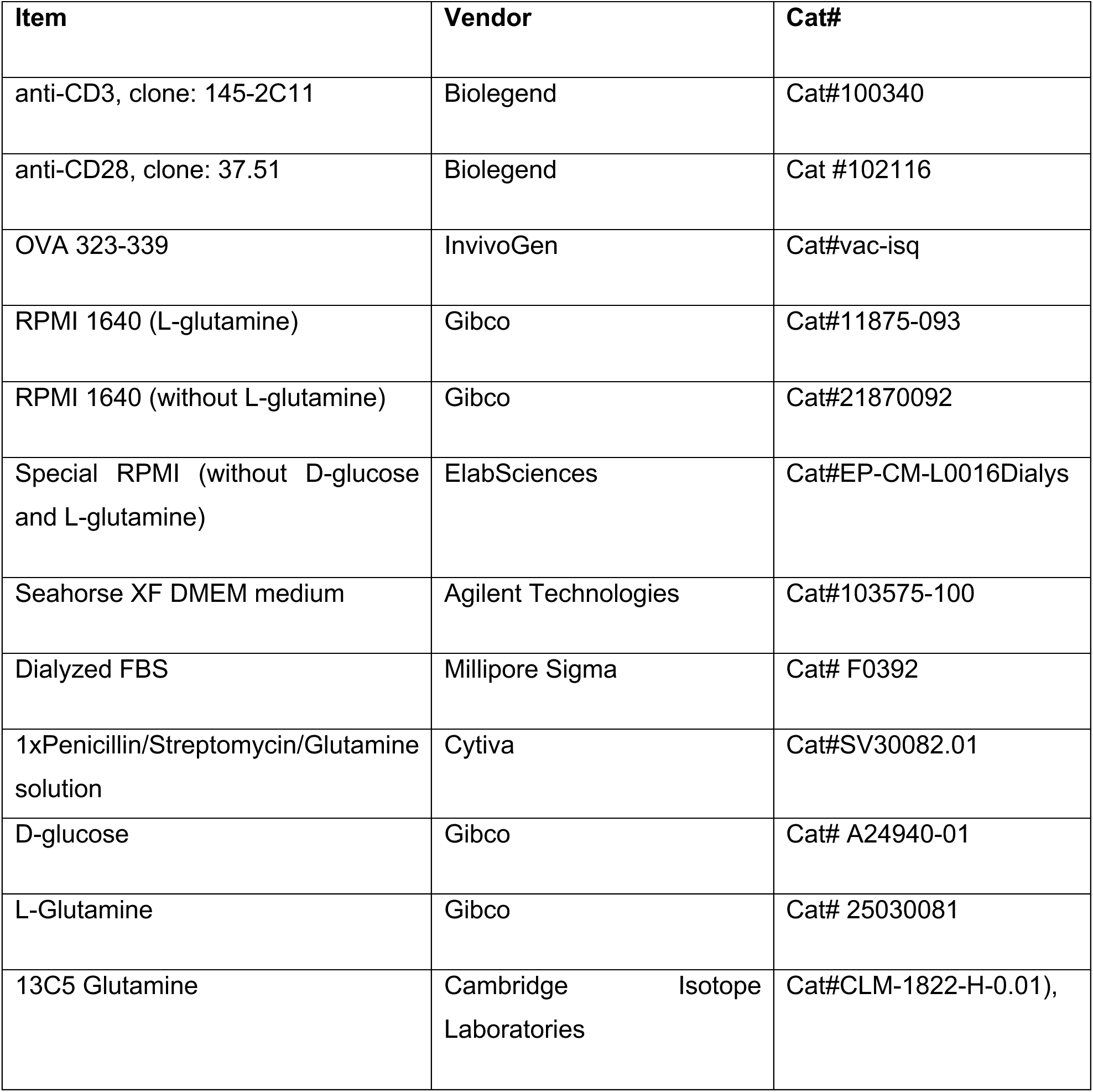
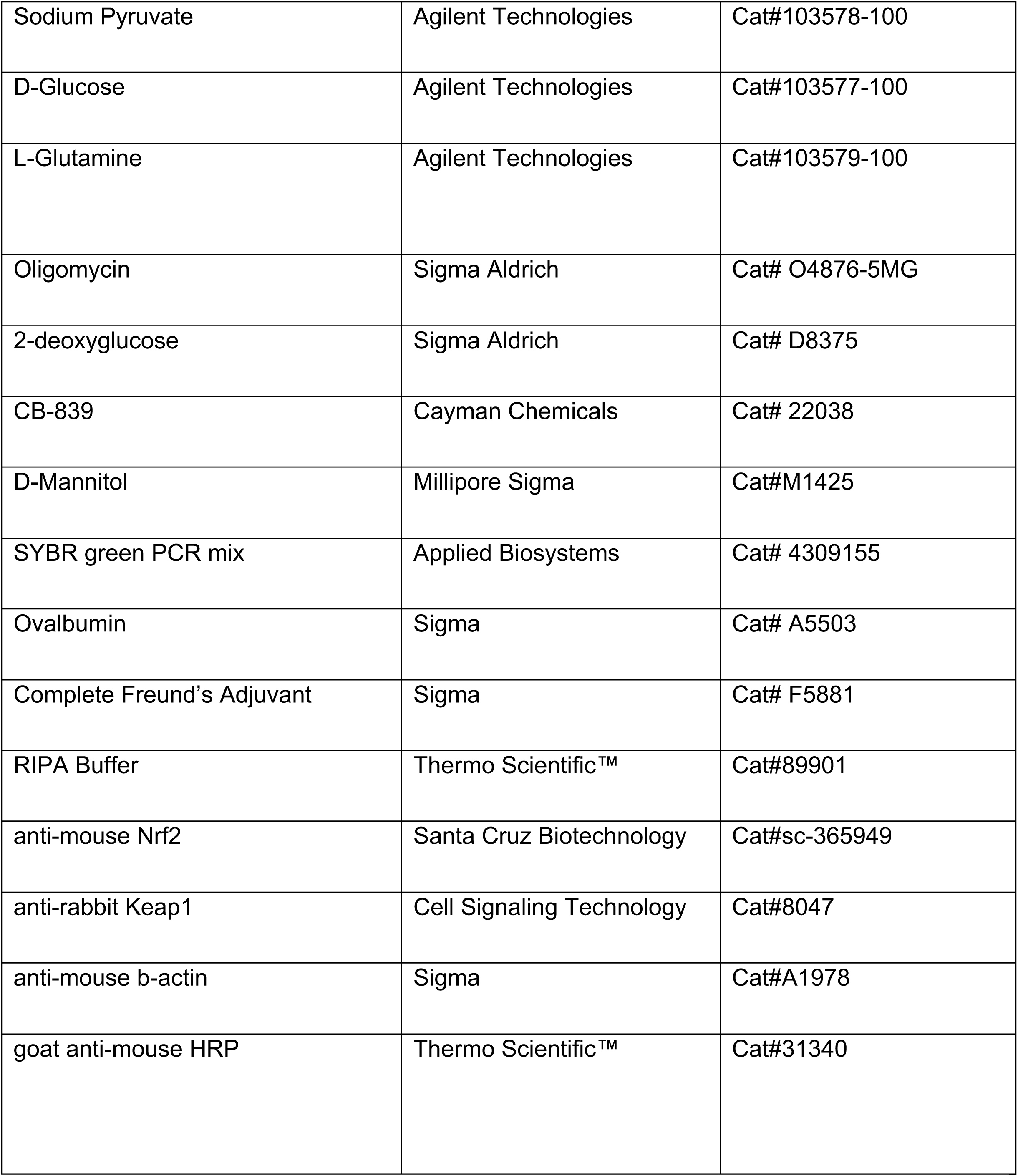
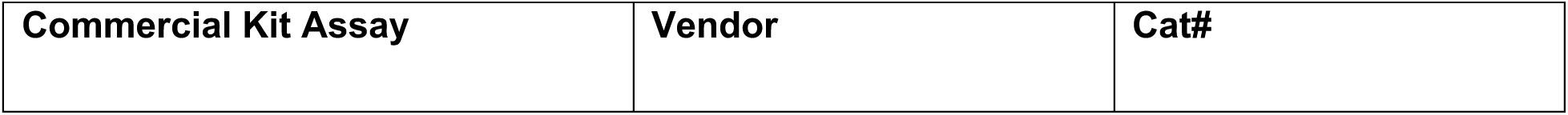
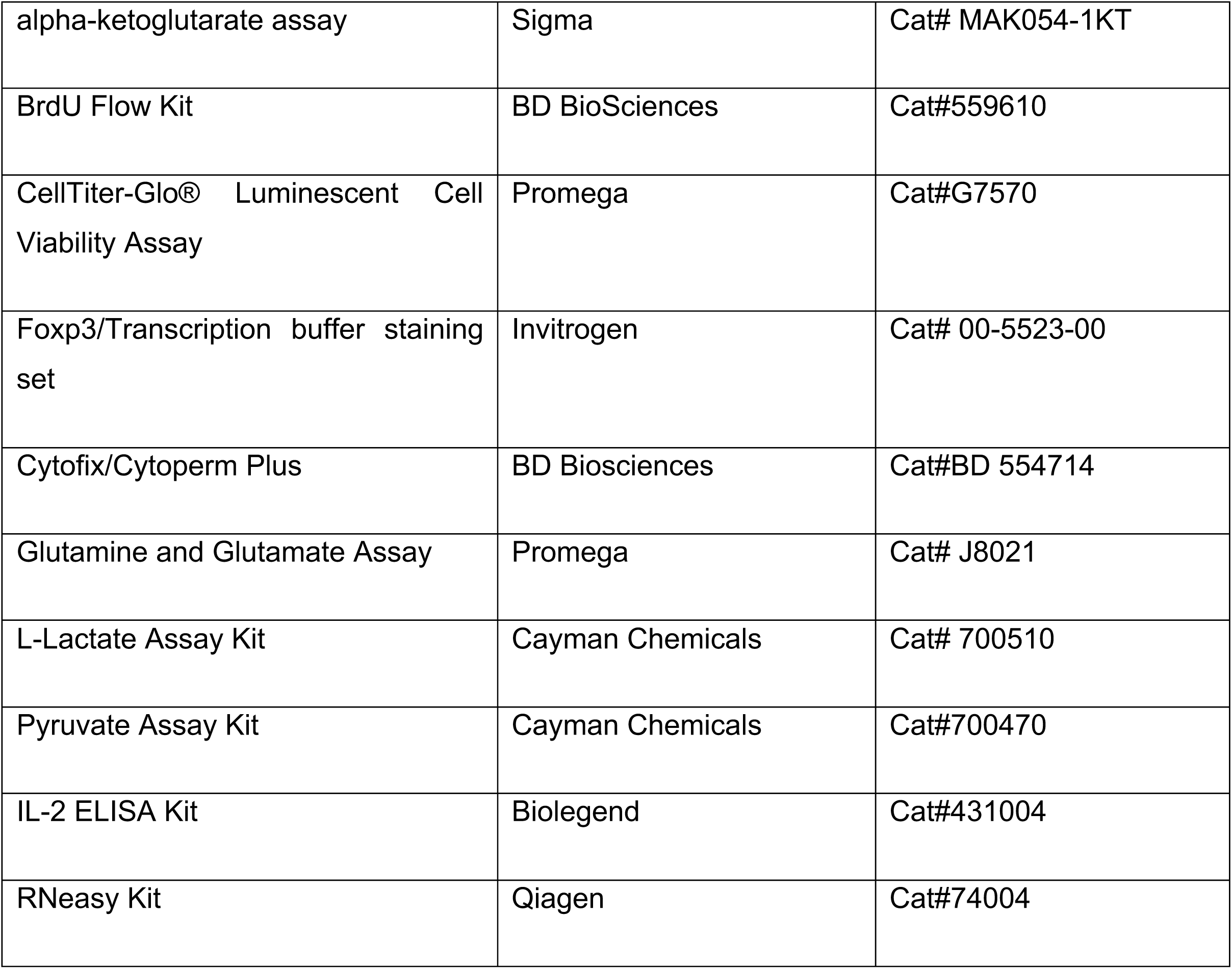

| Item | Vendor | Cat# |
| --- | --- | --- |
| anti-CD3, clone: 145-2C11 | Biolegend | Cat#100340 |
| anti-CD28, clone: 37.51 | Biolegend | Cat #102116 |
| OVA 323-339 | InvivoGen | Cat#vac-isq |
| RPMI 1640 (L-glutamine) | Gibco | Cat#11875-093 |
| RPMI 1640 (without L-glutamine) | Gibco | Cat#21870092 |
| Special RPMI (without D-glucose and L-glutamine) | ElabSciences | Cat#EP-CM-L0016Dialys |
| Seahorse XF DMEM medium | Agilent Technologies | Cat#103575-100 |
| Dialyzed FBS | Millipore Sigma | Cat# F0392 |
| 1xPenicillin/Streptomycin/Glutamine solution | Cytiva | Cat#SV30082.01 |
| D-glucose | Gibco | Cat# A24940-01 |
| L-Glutamine | Gibco | Cat# 25030081 |
| 13C5 Glutamine | Cambridge Isotope Laboratories | Cat#CLM-1822-H-0.01), |
| Sodium Pyruvate | Agilent Technologies | Cat#103578-100 |
| D-Glucose | Agilent Technologies | Cat#103577-100 |
| L-Glutamine | Agilent Technologies | Cat#103579-100 |
| Oligomycin | Sigma Aldrich | Cat# O4876-5MG |
| 2-deoxyglucose | Sigma Aldrich | Cat# D8375 |
| CB-839 | Cayman Chemicals | Cat# 22038 |
| D-Mannitol | Millipore Sigma | Cat#M1425 |
| SYBR green PCR mix | Applied Biosystems | Cat# 4309155 |
| Ovalbumin | Sigma | Cat# A5503 |
| Complete Freund's Adjuvant | Sigma | Cat# F5881 |
| RIPA Buffer | Thermo Scientific™ | Cat#89901 |
| anti-mouse Nrf2 | Santa Cruz Biotechnology | Cat#sc-365949 |
| anti-rabbit Keap1 | Cell Signaling Technology | Cat#8047 |
| anti-mouse b-actin | Sigma | Cat#A1978 |
| goat anti-mouse HRP | Thermo Scientific™ | Cat#31340 |

| Commercial Kit Assay | Vendor | Cat# |
| --- | --- | --- |
| alpha-ketoglutarate assay | Sigma | Cat# MAK054-1KT |
| BrdU Flow Kit | BD BioSciences | Cat#559610 |
| CellTiter-Glo® Luminescent Cell Viability Assay | Promega | Cat#G7570 |
| Foxp3/Transcription buffer staining set | Invitrogen | Cat# 00-5523-00 |
| Cytofix/Cytoperm Plus | BD Biosciences | Cat#BD 554714 |
| Glutamine and Glutamate Assay | Promega | Cat# J8021 |
| L-Lactate Assay Kit | Cayman Chemicals | Cat# 700510 |
| Pyruvate Assay Kit | Cayman Chemicals | Cat#700470 |
| IL-2 ELISA Kit | Biolegend | Cat#431004 |
| RNeasy Kit | Qiagen | Cat#74004 |

#### Oligonucleotides

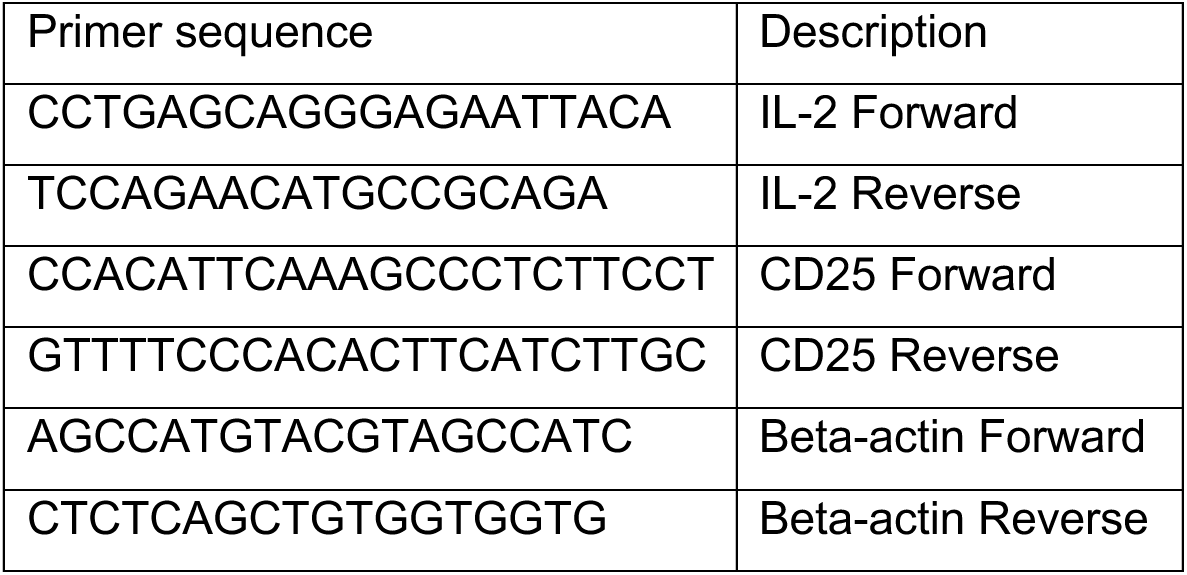

#### Software and Algorithm

|  |  |  |
| --- | --- | --- |
| bioRender | bioRender | http://www.biorender.com/ |
| FloJo ver. 10.8.2 | TreeStar software | https://www.flowjo.com/ |
| ImageJ | ImageJ | https://imagej.nih.gov/ij/ |
| GraphPad Prism ver. 10.0.0 | GraphPad software | http://www.graphpad.com/ |

---

**Supplemental Figure1.**
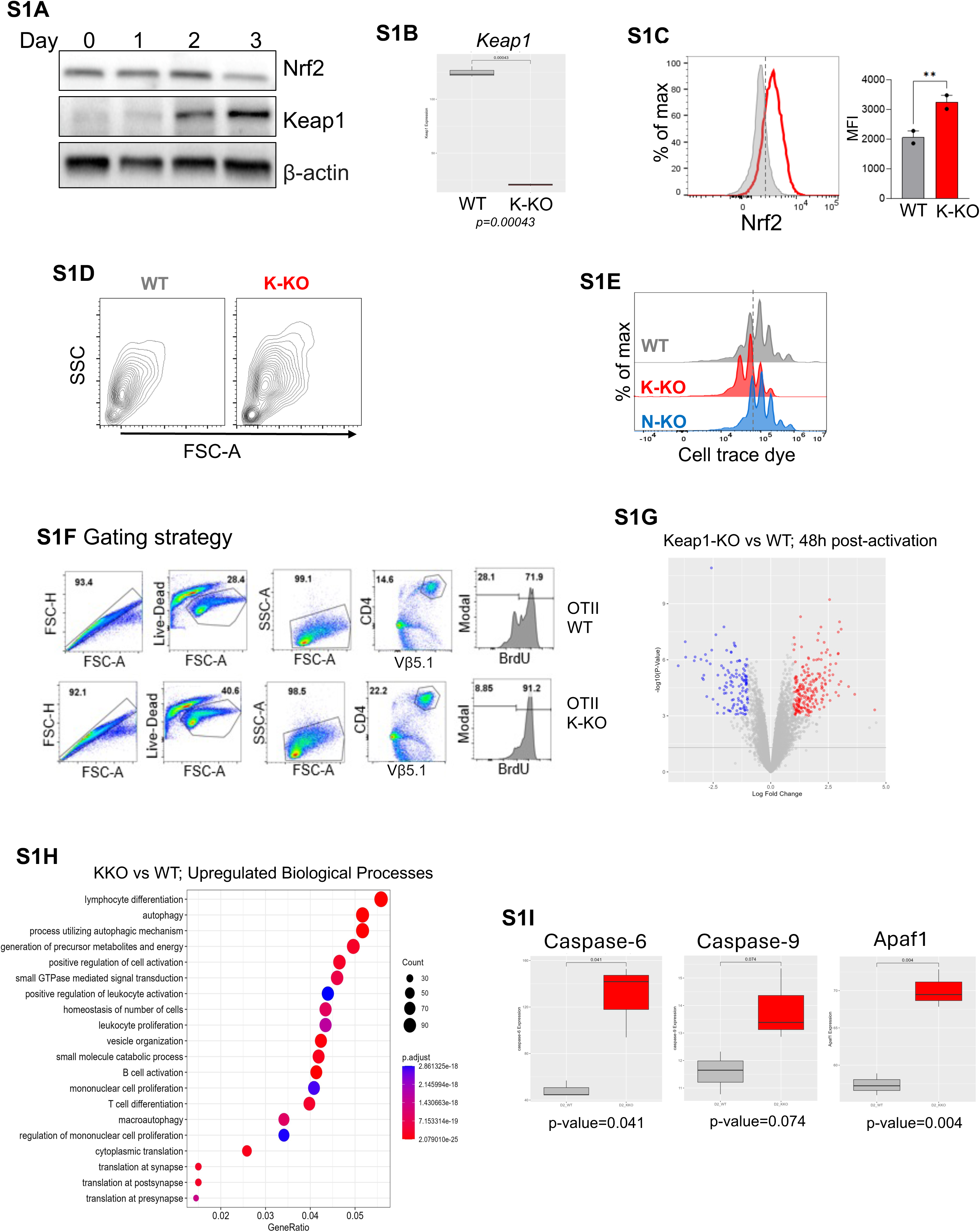
(A) CD4 T-cells isolated from C57BL/6 mice were stimulated using anti-CD3 and anti-CD28 antibodies. Immunoblot shows Nrf2 and Keap1 protein expression in the lysates of cells harvested after the indicated number of days. β-actin is shown as the loading control. Data is representative of two independent experiments. (B) Box plot comparing the mRNA levels of *Keap1* from Keap1-KO vs WT CD4 T-cells 48 h post-activation. Data are obtained from RNAseq analysis (n=3). (C) Representative histogram overlay and summary graph show Nrf2 protein expression in CD4 T-cells isolated from WT and Keap1-KO mice on day 2 after injection with anti-CD3 antibody intraperitoneally (n=2). (D) Flow cytometric plot showing the FSC and SSC of CD4 T-cells activated *in vitr*o. Data is representative of three independent experiments. (E) Representative histogram overlay showing proliferation of WT, Keap1-KO, and Nrf2-KO CD4 T-cells stimulated *in vitro* for 72 hours. Data is representative of three different experiments. (F) A representative gating strategy of flow cytometric analysis of data shown in Figure 1G. (G) Volcano plot showing the significantly upregulated (red) and downregulated (blue) genes identified from RNA-seq analysis of Keap1-KO vs WT CD4 T-cells 48 h post-activation. (n=3). (H) Gene Ontology data showing the upregulated genes involved in the biological processes in Keap1-KO over WT 48 h post-activation. Data are obtained from RNA-seq analysis (n=3). (I) Box plot comparing the mRNA levels of the indicated genes between Keap1-KO vs WT CD4 T-cells 48 h post-activation. Data are obtained from RNAseq analysis. Data are shown as mean ± SEM from the indicated number of sets of mice. *p<0.05, **p<0.01, ****p<0.0001

**Supplemental Figure 2.**
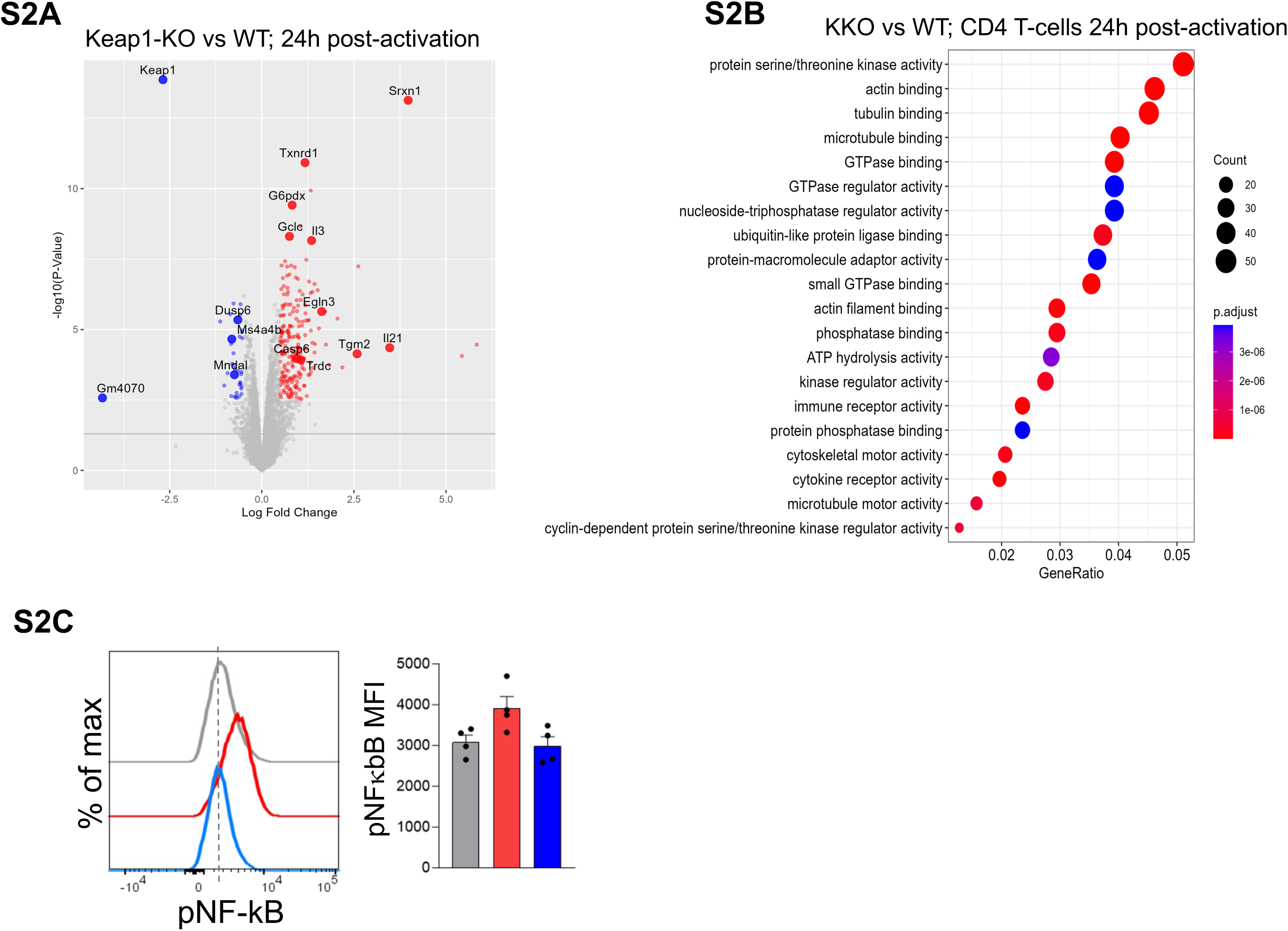
(A) Volcano plot showing the significantly upregulated (red) and downregulated (blue) genes in Keap1-KO vs WT CD4 T-cells 24h post-activation identified from RNA-seq analysis. (n=3). (B) Gene Ontology enrichment analysis of molecular functions for upregulated genes between Keap1KO vs WT CD4 T-cells 48h post activation. Data are obtained from RNA-seq analysis (n=3). (C) Representative histogram overlay and summary graph show the levels of phosphorylated NF-kB in WT, Keap1-KO, and Nrf2-KO CD4 T-cells activated in vitro for 24 h (n=4) Data are shown as mean ± SEM from the indicated number of sets of mice.

**Supplemental Figure 3.**
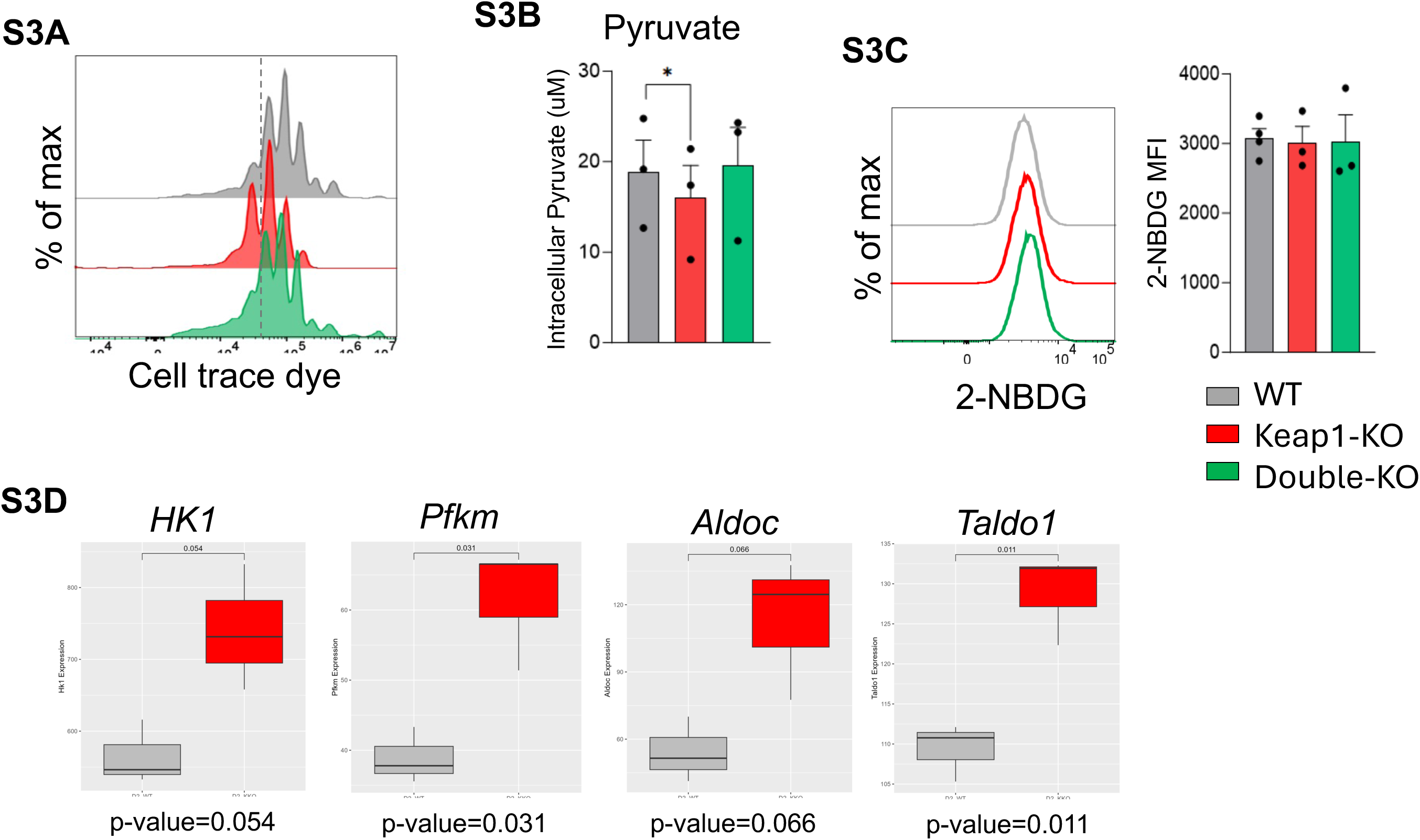
(A) Representative histogram overlay of the proliferation of WT, Keap1-KO, and Double-KO mice. Data is representative of three different experiments. (B) Intracellular levels of pyruvate were compared between WT, Keap1-KO, and Double-KO mice CD4 T-cells activated in vitro for 48 h (n=3). (C) Glucose uptake of WT, Keap1-KO, and Double-KO mice CD4 T-cells was measured 48h after activation by incubating with 2-NBDG for 30 mins followed by flow cytometry. Representative histogram overlay and summary graph are shown (n=3). (D) Box plots from RNAseq analysis comparing the mRNA levels of the indicated genes between Keap1-KO vs WT CD4 T-cells 48 h post-activation. Data are shown as mean ± SEM from the indicated number of sets of mice.

**Supplemental Figure 4.**
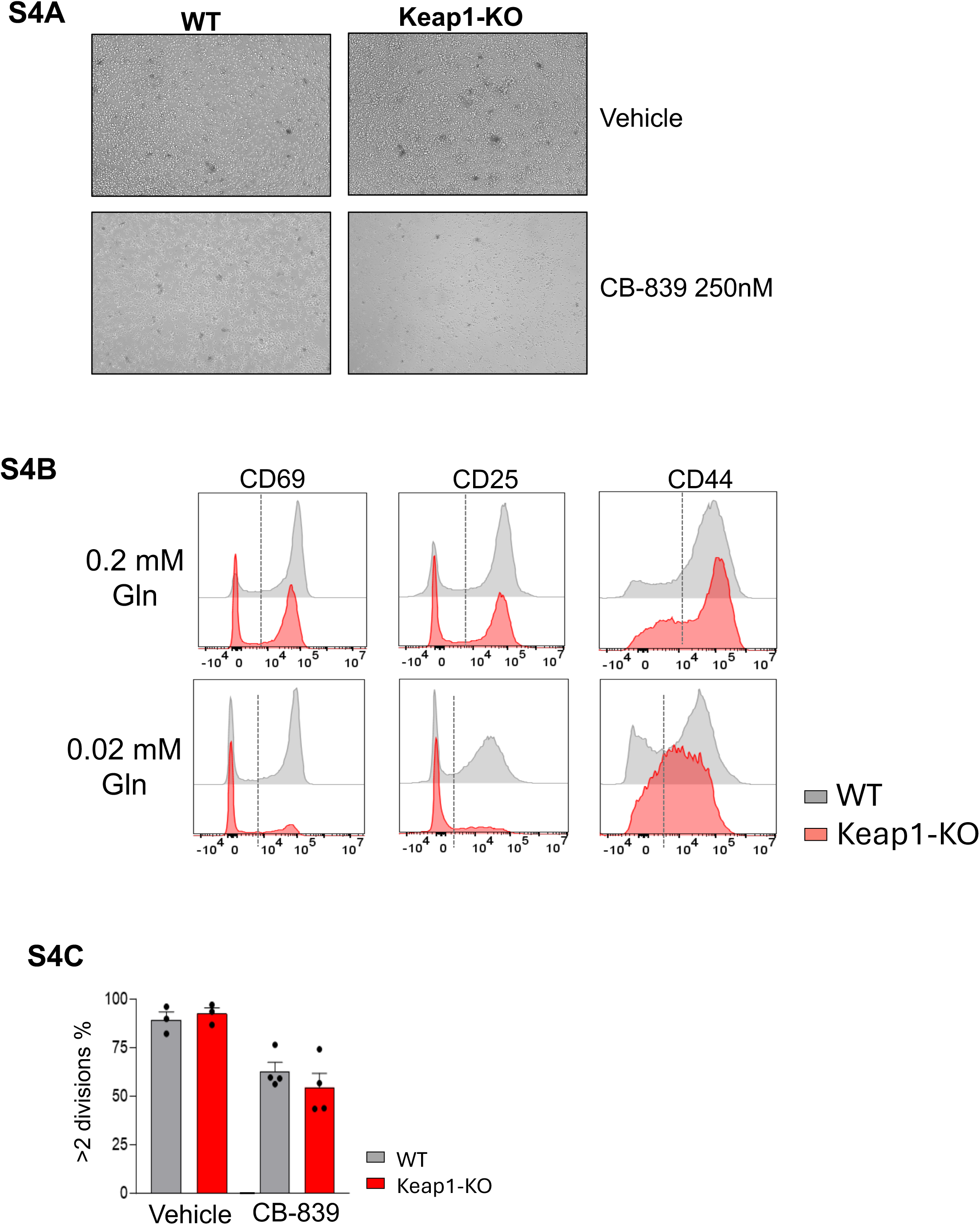
(A) Microscopic Images demonstrate the differences in cell morphology between WT and Keap1-KO CD4 T-cells activated *in vitro* for 96h in the presence of vehicle or CB-839. Data are representative of three independent experiments. (B) Representative histogram overlays showing the expression of CD25, CD44, and CD69 in WT and Keap1-KO CD4 T-cells activated *in vitro* in media with limiting glutamine (Gln). Data is representative of three independent experiments. (C) Summary graph showing cell proliferation of WT and Keap1-KO CD4 T-cells treated with Vehicle or CB-839 (n=3). Data are shown as mean ± SEM from the indicated number of sets of mice.

## References

1. Mondino, A., Khoruts, A., and Jenkins, M.K. (1996). The anatomy of T-cell activation and tolerance. Proc. Natl. Acad. Sci. U. S. A. 93, 2245–2252. 10.1073/pnas.93.6.2245.

2. Smith-Garvin, J.E., Koretzky, G.A., and Jordan, M.S. (2009). T cell activation. Annu. Rev. Immunol. 27, 591–619. 10.1146/annurev.immunol.021908.132706.

3. Jenkins, M.K., Khoruts, A., Ingulli, E., Mueller, D.L., McSorley, S.J., Reinhardt, R.L., Itano, A., and Pape, K.A. (2001). In vivo activation of antigen-specific CD4 T cells. Annu. Rev. Immunol. 19, 23–45. 10.1146/annurev.immunol.19.1.23.

4. Shyer, J.A., Flavell, R.A., and Bailis, W. (2020). Metabolic signaling in T cells. Cell Res. 30, 649–659. 10.1038/s41422-020-0379-5.

5. Buck, M.D., O’Sullivan, D., and Pearce, E.L. (2015). T cell metabolism drives immunity. J. Exp. Med. 212, 1345–1360. 10.1084/jem.20151159.

6. Pearce, E.L. (2010). Metabolism in T cell activation and differentiation. Curr. Opin. Immunol. 22, 314–320. 10.1016/j.coi.2010.01.018.

7. Pearce, E.L., Poffenberger, M.C., Chang, C.-H., and Jones, R.G. (2013). Fueling Immunity: Insights into Metabolism and Lymphocyte Function. Science 342, 1242454. 10.1126/science.1242454.

8. Almeida, L., Lochner, M., Berod, L., and Sparwasser, T. (2016). Metabolic pathways in T cell activation and lineage differentiation. Semin. Immunol. 28, 514–524. 10.1016/j.smim.2016.10.009.

9. Carr, E.L., Kelman, A., Wu, G.S., Gopaul, R., Senkevitch, E., Aghvanyan, A., Turay, A.M., and Frauwirth, K.A. (2010). Glutamine Uptake and Metabolism Are Coordinately Regulated by ERK/MAPK During T Lymphocyte Activation. J. Immunol. Baltim. Md 1950 *185*, 1037– 1044. 10.4049/jimmunol.0903586.

10. Nakaya, M., Xiao, Y., Zhou, X., Chang, J.-H., Chang, M., Cheng, X., Blonska, M., Lin, X., and Sun, S.-C. (2014). Inflammatory T Cell Responses Rely on Amino Acid Transporter ASCT2 Facilitation of Glutamine Uptake and mTORC1 Kinase Activation. Immunity 40, 692–705. 10.1016/j.immuni.2014.04.007.

11. Sinclair, L.V., Rolf, J., Emslie, E., Shi, Y.-B., Taylor, P.M., and Cantrell, D.A. (2013). Control of amino-acid transport by antigen receptors coordinates the metabolic reprogramming essential for T cell differentiation. Nat. Immunol. 14, 500–508. 10.1038/ni.2556.

12. Murphy, M.P., and Siegel, R.M. (2013). Mitochondrial ROS Fire Up T Cell Activation. Immunity 38, 201–202. 10.1016/j.immuni.2013.02.005.

13. Lepez, A., Pirnay, T., Denanglaire, S., Perez-Morga, D., Vermeersch, M., Leo, O., and Andris, F. (2020). Long-term T cell fitness and proliferation is driven by AMPK-dependent regulation of reactive oxygen species. Sci. Rep. 10, 21673. 10.1038/s41598-020-78715-2.

14. Yarosz, E.L., and Chang, C.-H. (2018). The Role of Reactive Oxygen Species in Regulating T Cell-mediated Immunity and Disease. Immune Netw. 18, e14. 10.4110/in.2018.18.e14.

15. Franchina, D.G., Dostert, C., and Brenner, D. (2018). Reactive oxygen species: involvement in T cell signaling and metabolism. Trends Immunol. 39, 489–502.

16. Kesarwani, P., Murali, A.K., Al-Khami, A.A., and Mehrotra, S. (2013). Redox Regulation of T-Cell Function: From Molecular Mechanisms to Significance in Human Health and Disease. Antioxid. Redox Signal. 18, 1497–1534. 10.1089/ars.2011.4073.

17. Peng, H.-Y., Lucavs, J., Ballard, D., Das, J.K., Kumar, A., Wang, L., Ren, Y., Xiong, X., and Song, J. (2021). Metabolic Reprogramming and Reactive Oxygen Species in T Cell Immunity. Front. Immunol. 12, 652687. 10.3389/fimmu.2021.652687.

18. Itoh, K., Wakabayashi, N., Katoh, Y., Ishii, T., Igarashi, K., Engel, J.D., and Yamamoto, M. (1999). Keap1 represses nuclear activation of antioxidant responsive elements by Nrf2 through binding to the amino-terminal Neh2 domain. Genes Dev. 13, 76–86. 10.1101/gad.13.1.76.

19. Deshmukh, P., Unni, S., Krishnappa, G., and Padmanabhan, B. (2017). The Keap1-Nrf2 pathway: promising therapeutic target to counteract ROS-mediated damage in cancers and neurodegenerative diseases. Biophys. Rev. 9, 41–56. 10.1007/s12551-016-0244-4.

20. Baird, L., and Yamamoto, M. (2020). The Molecular Mechanisms Regulating the KEAP1-NRF2 Pathway. Mol. Cell. Biol. 40, e00099–20. 10.1128/MCB.00099-20.

21. Taguchi, K., Motohashi, H., and Yamamoto, M. (2011). Molecular mechanisms of the Keap1–Nrf2 pathway in stress response and cancer evolution. Genes Cells 16, 123–140. 10.1111/j.1365-2443.2010.01473.x.

22. He, F., Ru, X., and Wen, T. (2020). NRF2, a Transcription Factor for Stress Response and Beyond. Int. J. Mol. Sci. 21, 4777. 10.3390/ijms21134777.

23. Xu, Y., Yu, Z., Fu, H., Guo, Y., Hu, P., and Shi, J. (2022). Dual Inhibitions on Glucose/Glutamine Metabolisms for Nontoxic Pancreatic Cancer Therapy. ACS Appl. Mater. Interfaces 14, 21836–21847. 10.1021/acsami.2c00111.

24. Jin, J., Byun, J.-K., Choi, Y.-K., and Park, K.-G. (2023). Targeting glutamine metabolism as a therapeutic strategy for cancer. Exp. Mol. Med. 55, 706–715. 10.1038/s12276-023-00971-9.

25. Pyaram, K., Kumar, A., Kim, Y.-H., Noel, S., Reddy, S.P., Rabb, H., and Chang, C.-H. (2019). Keap1-Nrf2 System Plays an Important Role in Invariant Natural Killer T Cell Development and Homeostasis. Cell Rep. 27, 699–707.e4. 10.1016/j.celrep.2019.03.052.

26. Rajendiran, A., Subramanyam, S.H., Klemm, P., Jankowski, V., van Loosdregt, J., Vastert, B., Vollbach, K., Wagner, N., Tenbrock, K., and Ohl, K. (2022). NRF2/Itaconate Axis Regulates Metabolism and Inflammatory Properties of T Cells in Children with JIA. Antioxid. Basel Switz. 11, 2426. 10.3390/antiox11122426.

27. Gnanaprakasam, J.N.R., Kushwaha, B., Liu, L., Chen, X., Kang, S., Wang, T., Cassel, T.A., Adams, C.M., Higashi, R.M., Scott, D.A., et al. (2023). Asparagine restriction enhances CD8+ T cell metabolic fitness and antitumoral functionality through an NRF2-dependent stress response. Nat. Metab. 5, 1423–1439. 10.1038/s42255-023-00856-1.

28. Turley, A.E., Zagorski, J.W., and Rockwell, C.E. (2015). The Nrf2 activator tBHQ inhibits T cell activation of primary human CD4 T cells. Cytokine 71, 289–295. 10.1016/j.cyto.2014.11.006.

29. Zagorski, J.W., Turley, A.E., Freeborn, R.A., VanDenBerg, K.R., Dover, H.E., Kardell, B.R., Liby, K.T., and Rockwell, C.E. (2018). Differential effects of the Nrf2 activators tBHQ and CDDO-Im on the early events of T cell activation. Biochem. Pharmacol. 147, 67–76. 10.1016/j.bcp.2017.11.005.

30. Wu, Q., Wang, Q., Mao, G., Dowling, C.A., Lundy, S.K., and Mao-Draayer, Y. (2017). Dimethyl Fumarate Selectively Reduces Memory T Cells and Shifts the Balance between Th1/Th17 and Th2 in Multiple Sclerosis Patients. J. Immunol. Baltim. Md 1950 *198*, 3069– 3080. 10.4049/jimmunol.1601532.

31. Morzadec, C., Macoch, M., Sparfel, L., Kerdine-Römer, S., Fardel, O., and Vernhet, L. (2014). Nrf2 expression and activity in human T lymphocytes: stimulation by T cell receptor activation and priming by inorganic arsenic and tert-butylhydroquinone. Free Radic. Biol. Med. 71, 133–145. 10.1016/j.freeradbiomed.2014.03.006.

32. Noel, S., Martina, M.N., Bandapalle, S., Racusen, L.C., Potteti, H.R., Hamad, A.R.A., Reddy, S.P., and Rabb, H. (2015). T Lymphocyte-Specific Activation of Nrf2 Protects from AKI. J. Am. Soc. Nephrol. JASN 26, 2989–3000. 10.1681/ASN.2014100978.

33. Dimeloe, S., Burgener, A.-V., Grählert, J., and Hess, C. (2017). T-cell metabolism governing activation, proliferation and differentiation; a modular view. Immunology 150, 35–44. 10.1111/imm.12655.

34. Barnden, M.J., Allison, J., Heath, W.R., and Carbone, F.R. (1998). Defective TCR expression in transgenic mice constructed using cDNA-based alpha-and beta-chain genes under the control of heterologous regulatory elements. Immunol. Cell Biol. 76, 34–40. 10.1046/j.1440-1711.1998.00709.x.

35. Shah, K., Al-Haidari, A., Sun, J., and Kazi, J.U. (2021). T cell receptor (TCR) signaling in health and disease. Signal Transduct. Target. Ther. 6, 412. 10.1038/s41392-021-00823-w.

36. Au-Yeung, B.B., Zikherman, J., Mueller, J.L., Ashouri, J.F., Matloubian, M., Cheng, D.A., Chen, Y., Shokat, K.M., and Weiss, A. (2014). A sharp T-cell antigen receptor signaling threshold for T-cell proliferation. Proc. Natl. Acad. Sci. U. S. A. 111, E3679–3688. 10.1073/pnas.1413726111.

37. Hwang, J.-R., Byeon, Y., Kim, D., and Park, S.-G. (2020). Recent insights of T cell receptor-mediated signaling pathways for T cell activation and development. Exp. Mol. Med. 52, 750–761. 10.1038/s12276-020-0435-8.

38. Ashouri, J.F., and Weiss, A. (2017). Endogenous Nur77 Is a Specific Indicator of Antigen Receptor Signaling in Human T and B Cells. J. Immunol. Baltim. Md 1950 *198*, 657–668. 10.4049/jimmunol.1601301.

39. Wang, H., Kadlecek, T.A., Au-Yeung, B.B., Goodfellow, H.E.S., Hsu, L.-Y., Freedman, T.S., and Weiss, A. (2010). ZAP-70: an essential kinase in T-cell signaling. Cold Spring Harb. Perspect. Biol. 2, a002279. 10.1101/cshperspect.a002279.

40. Adachi, K., and Davis, M.M. (2011). T-cell receptor ligation induces distinct signaling pathways in naive vs. antigen-experienced T cells. Proc. Natl. Acad. Sci. U. S. A. 108, 1549–1554. 10.1073/pnas.1017340108.

41. Bachmann, M.F., and Oxenius, A. (2007). Interleukin 2: from immunostimulation to immunoregulation and back again. EMBO Rep. 8, 1142–1148. 10.1038/sj.embor.7401099.

42. Ross, S.H., and Cantrell, D.A. (2018). Signaling and Function of Interleukin-2 in T Lymphocytes. Annu. Rev. Immunol. 36, 411–433. 10.1146/annurev-immunol-042617-053352.

43. Jones, D.M., Read, K.A., and Oestreich, K.J. (2020). Dynamic Roles for IL-2-STAT5 Signaling in Effector and Regulatory CD4+ T Cell Populations. J. Immunol. Baltim. Md 1950 *205*, 1721–1730. 10.4049/jimmunol.2000612.

44. Waickman, A.T., and Powell, J.D. (2012). mTOR, metabolism, and the regulation of T-cell differentiation and function. Immunol. Rev. 249, 43–58. 10.1111/j.1600-065X.2012.01152.x.

45. Palmer, C.S., Ostrowski, M., Balderson, B., Christian, N., and Crowe, S.M. (2015). Glucose metabolism regulates T cell activation, differentiation, and functions. Front. Immunol. 6, 1. 10.3389/fimmu.2015.00001.

46. Li, X., Yang, Y., Zhang, B., Lin, X., Fu, X., An, Y., Zou, Y., Wang, J.-X., Wang, Z., and Yu, T. (2022). Lactate metabolism in human health and disease. Signal Transduct. Target. Ther. 7, 305. 10.1038/s41392-022-01151-3.

47. Rogatzki, M.J., Ferguson, B.S., Goodwin, M.L., and Gladden, L.B. (2015). Lactate is always the end product of glycolysis. Front. Neurosci. 9, 22. 10.3389/fnins.2015.00022.

48. Wapnir, R.A., and Stiel, L. (1985). Regulation of gluconeogenesis by glycerol and its phosphorylated derivatives. Biochem. Med. 33, 141–148. 10.1016/0006-2944(85)90022-5.

49. Song, J., Sun, H., Zhang, S., and Shan, C. (2022). The Multiple Roles of Glucose-6-Phosphate Dehydrogenase in Tumorigenesis and Cancer Chemoresistance. Life 12, 271. 10.3390/life12020271.

50. Ma, G., Zhang, Z., Li, P., Zhang, Z., Zeng, M., Liang, Z., Li, D., Wang, L., Chen, Y., Liang, Y., et al. (2022). Reprogramming of glutamine metabolism and its impact on immune response in the tumor microenvironment. Cell Commun. Signal. 20, 114. 10.1186/s12964-022-00909-0.

51. Hörig, H., Spagnoli, G.C., Filgueira, L., Babst, R., Gallati, H., Harder, F., Juretic, A., and Heberer, M. (1993). Exogenous glutamine requirement is confined to late events of T cell activation. J. Cell. Biochem. 53, 343–351. 10.1002/jcb.240530412.

52. Mu, P., Huo, J., Li, X., Li, W., Li, X., Ao, J., and Chen, X. (2022). IL-2 Signaling Couples the MAPK and mTORC1 Axes to Promote T Cell Proliferation and Differentiation in Teleosts. J. Immunol. Baltim. Md 1950 *208*, 1616–1631. 10.4049/jimmunol.2100764.

53. Mitsuishi, Y., Taguchi, K., Kawatani, Y., Shibata, T., Nukiwa, T., Aburatani, H., Yamamoto, M., and Motohashi, H. (2012). Nrf2 redirects glucose and glutamine into anabolic pathways in metabolic reprogramming. Cancer Cell 22, 66–79. 10.1016/j.ccr.2012.05.016.

54. He, F., Antonucci, L., and Karin, M. (2020). NRF2 as a regulator of cell metabolism and inflammation in cancer. Carcinogenesis 41, 405–416. 10.1093/carcin/bgaa039.

55. Zhang, J., Pavlova, N.N., and Thompson, C.B. (2017). Cancer cell metabolism: the essential role of the nonessential amino acid, glutamine. EMBO J. 36, 1302–1315. 10.15252/embj.201696151.

56. Daye, D., and Wellen, K.E. (2012). Metabolic reprogramming in cancer: unraveling the role of glutamine in tumorigenesis. Semin. Cell Dev. Biol. 23, 362–369. 10.1016/j.semcdb.2012.02.002.

57. Hui, S., Cowan, A.J., Zeng, X., Yang, L., TeSlaa, T., Li, X., Bartman, C., Zhang, Z., Jang, C., Wang, L., et al. (2020). Quantitative Fluxomics of Circulating Metabolites. Cell Metab. 32, 676–688.e4. 10.1016/j.cmet.2020.07.013.

58. Iwao, H., Fukui, K., Yamamoto, A., Shoji, T., and Abe, Y. (1987). Effects of captopril on plasma atrial natriuretic polypeptide and the renin angiotensin system in rats. Clin. Exp. Hypertens. A. 9, 697–701. 10.3109/10641968709164245.

59. Varghese, S., Pramanik, S., Williams, L.J., Hodges, H.R., Hudgens, C.W., Fischer, G.M., Luo, C.K., Knighton, B., Tan, L., Lorenzi, P.L., et al. (2021). The Glutaminase Inhibitor CB-839 (Telaglenastat) Enhances the Antimelanoma Activity of T-Cell-Mediated Immunotherapies. Mol. Cancer Ther. 20, 500–511. 10.1158/1535-7163.MCT-20-0430.

60. Pant, A., Dasgupta, D., Tripathi, A., and Pyaram, K. (2023). Beyond Antioxidation: Keap1-Nrf2 in the Development and Effector Functions of Adaptive Immune Cells. ImmunoHorizons 7, 288–298. 10.4049/immunohorizons.2200061.

61. Moriggl, R., Topham, D.J., Teglund, S., Sexl, V., McKay, C., Wang, D., Hoffmeyer, A., van Deursen, J., Sangster, M.Y., Bunting, K.D., et al. (1999). Stat5 Is Required for IL-2-Induced Cell Cycle Progression of Peripheral T Cells. Immunity 10, 249–259. 10.1016/S1074-7613(00)80025-4.

62. Finlay, D.K. (2012). Regulation of glucose metabolism in T cells: new insight into the role of Phosphoinositide 3-kinases. Front. Immunol. 3. 10.3389/fimmu.2012.00247.

63. Chang, C.-H., Curtis, J.D., Maggi, L.B., Faubert, B., Villarino, A.V., O’Sullivan, D., Huang, S.C.-C., Van Der Windt, G.J., Blagih, J., and Qiu, J. (2013). Posttranscriptional control of T cell effector function by aerobic glycolysis. Cell 153, 1239–1251.

64. Myers, D.R., Wheeler, B., and Roose, J.P. (2019). mTOR and Other Effector Kinase Signals that Impact T Cell Function and Activity. Immunol. Rev. 291, 134–153. 10.1111/imr.12796.

65. Wu, K.C., Cui, J.Y., and Klaassen, C.D. (2011). Beneficial role of Nrf2 in regulating NADPH generation and consumption. Toxicol. Sci. Off. J. Soc. Toxicol. 123, 590–600. 10.1093/toxsci/kfr183.

66. Crawford, J., and Cohen, H.J. (1985). The essential role of L-glutamine in lymphocyte differentiation in vitro. J. Cell. Physiol. 124, 275–282. 10.1002/jcp.1041240216.

67. Yaqoob, P., and Calder, P.C. (1997). Glutamine requirement of proliferating T lymphocytes. Nutrition 13, 646–651. 10.1016/S0899-9007(97)83008-0.

68. Tajima, M., and Strober, W. (2022). Evaluation of Glutaminolysis in T cells. Curr. Protoc. 2, e540. 10.1002/cpz1.540.

69. Li, S., Takasu, C., Lau, H., Robles, L., Vo, K., Farzaneh, T., Vaziri, N.D., Stamos, M.J., and Ichii, H. (2020). Dimethyl Fumarate Alleviates Dextran Sulfate Sodium-Induced Colitis, through the Activation of Nrf2-Mediated Antioxidant and Anti-inflammatory Pathways. Antioxidants 9, 354. 10.3390/antiox9040354.

70. Ma, C., Gu, C., Lian, P., Wazir, J., Lu, R., Ruan, B., Wei, L., Li, L., Pu, W., Peng, Z., et al. (2023). Sulforaphane alleviates psoriasis by enhancing antioxidant defense through KEAP1-NRF2 Pathway activation and attenuating inflammatory signaling. Cell Death Dis. 14, 1–12. 10.1038/s41419-023-06234-9.

71. Mantione, M.E., Meloni, M., Sana, I., Bordini, J., Del Nero, M., Riba, M., Ranghetti, P., Perotta, E., Ghia, P., Scarfò, L., et al. (2024). Disrupting pro-survival and inflammatory pathways with dimethyl fumarate sensitizes chronic lymphocytic leukemia to cell death. Cell Death Dis. 15, 1–13. 10.1038/s41419-024-06602-z.

72. Reddy, N.M., Potteti, H.R., Mariani, T.J., Biswal, S., and Reddy, S.P. (2011). Conditional deletion of Nrf2 in airway epithelium exacerbates acute lung injury and impairs the resolution of inflammation. Am. J. Respir. Cell Mol. Biol. 45, 1161–1168. 10.1165/rcmb.2011-0144OC.

73. Babraham Bioinformatics-FastQC A Quality Control tool for High Throughput Sequence Data https://www.bioinformatics.babraham.ac.uk/projects/fastqc/.

74. RSEM: accurate transcript quantification from RNA-Seq data with or without a reference genome | BMC Bioinformatics | Full Text https://bmcbioinformatics.biomedcentral.com/articles/10.1186/1471-2105-12-323.

75. Robinson, M.D., McCarthy, D.J., and Smyth, G.K. (2010). edgeR: a Bioconductor package for differential expression analysis of digital gene expression data. Bioinforma. Oxf. Engl. 26, 139–140. 10.1093/bioinformatics/btp616.

76. Yu, G., Wang, L.-G., Han, Y., and He, Q.-Y. (2012). clusterProfiler: an R package for comparing biological themes among gene clusters. Omics J. Integr. Biol. 16, 284–287. 10.1089/omi.2011.0118.

77. ggplot2: Elegant Graphics for Data Analysis by WICKHAM, H. | Biometrics | Oxford Academic https://academic.oup.com/biometrics/article/67/2/678/7381027.

